# A Minimal Model of CD95 Signal Initiation Revealed by Advanced Super-resolution and Multiparametric Fluorescence Microscopy

**DOI:** 10.1101/2022.11.29.518370

**Authors:** Nina Bartels, Nicolaas T M van der Voort, Annemarie Greife, Arthur Bister, Constanze Wiek, Claus A M Seidel, Cornelia Monzel

**Author notes:** contributed equally (Nina Bartels, Nicolaas T M van der Voort).

## Abstract

Unraveling the spatiotemporal organization and dynamical interactions of receptors in the plasma membrane remains a key challenge for our mechanistic understanding of cell signal initiation. A paradigm of such process is the oligomerization of TNF receptor CD95 during apoptosis signaling, where molecular configurations are yet to be defined. Here, we scrutinize proposed oligomerization models in live cells, establishing a molecular sensitive imaging toolkit including time-resolved FRET spectroscopy, quantitative STED microscopy, confocal Photobleaching Step Analysis and FCS. CD95 interactions were probed over molecular concentrations of few to ∼ 1000 molecules/µm^2^, over ns to hours, and molecular to cellular scales. We further established high-fidelity monomer and dimer controls for quantitative benchmarking. Efficient apoptosis was already observed when ∼ 8 to 17% monomeric CD95 oligomerize to dimers/trimers after ligand binding. Our multiscale study highlights the importance of molecular concentrations, of the native environment, and reveals a minimal oligomerization model of CD95 signal initiation.

## Introduction

Identifying the spatiotemporal organization and dynamical interactions of receptors in the plasma membrane is key to our understanding of cell signal initiation. So far, we know about the molecules participating in distinct signaling cascades, however, insights about interaction networks, assembly kinetics, the formation of supramolecular patterns, as well as the role of molecular concentration remain sparse.

A paradigm of signal initiation is given by the characteristic molecular organization proposed for tumor necrosis factor receptors (TNFR), with the most prominent molecular configurations described below. The understanding of TNFR induced signaling is important, as these receptors initiate signaling for cell proliferation, morphogenesis and most prominently, cell apoptosis^1-3^. TNFRs are further targets of therapeutic approaches for various diseases, including cancer, autoimmunity, or infectious diseases^4^. Of particular interest is, in this context, the TNF receptor Cluster of Differentiation 95 (CD95/ Fas / TNFR6), as it is exclusively activated by the trimeric ligand CD95L (FasL / TNFL6 / CD178), thus providing high control over the stimulation of the receptor (Figure 1a).

**Figure 1:**
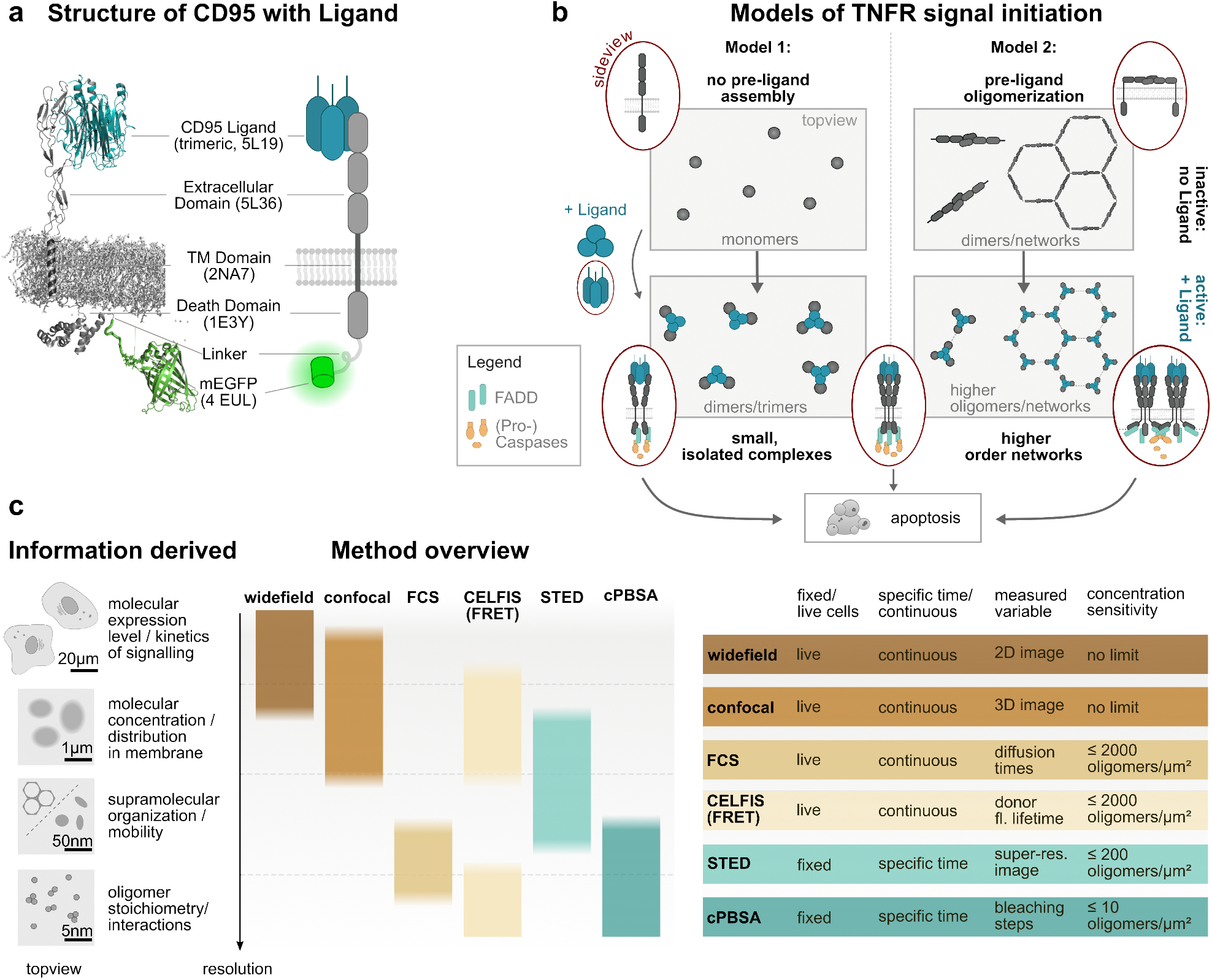
Probing Cluster of Differentiation 95 (CD95) signal initiation over a broad range of molecular concentrations and in space and time. **a)** Molecular structure and cartoon of CD95 receptor with genetically fused mEGFP and trimeric CD95 Ligand (CD95L). The four letter abbreviations refer to Protein Data Bank IDs. **b)** Schematic illustration of two proposed models of TNFR signal initiation. Left: monomeric receptors bind trimeric TNF ligands and form up to trimer-trimer receptor-ligand configurations. In the receptor activated state, the intracellular death domain (DD) opens and allows for recruitment of an adaptor molecule. The adapter molecule in case of CD95 is Fas-associated death domain protein (FADD, indicated in the sideview cartoons). A cascade of (Pro-)caspase binding and activation is initiated thereafter, resulting in intracellular protein cleavage followed by cell apoptosis. Right: TNFRs form inactive dimers prior to activation, which in turn assemble into a supramolecular honeycomb lattice consisting of hexagonal units of ∼ 24 nm diameter (sizes may vary with TNF receptor)^9^. After ligand binding to the lattice, the receptor dimers decouple, turn into their active state, and recruit FADD to the opened DDs. In the following, FADD may crosslink the DDs to reestablish the honeycomb lattice on the intracellular membrane side from which the (Pro-)caspase cascade evolves as in Model 1. **c)** Overview of test strategy using a combination of super-resolution and multiparametric fluorescence microscopy techniques covering cellular to single molecule scales. Fixed cell analyses at specific time-points are complemented by live cell studies over several hours. A concentration range spanning few to several 1000 molecules / µm^2^ is probed. Next to regular widefield, confocal time-lapse microscopy and FCS to monitor receptor and cell apoptosis dynamics, information about CD95 interaction dynamics, molecular distribution, and stoichiometries is obtained via Cell Lifetime FRET Image Spectroscopy (CELFIS), quantitative spot analysis using STED, and confocal Photobleaching Step Analysis (cPBSA). For further details see text and Methods.

Two models of TNFR oligomerization are primarily discussed to explain the molecular mechanisms underlying signal initiation (Figure 1b)^5-8^: the first model proposes initially monomeric receptors which, upon binding of the trimeric TNF ligand, recruit further receptors to form small signaling units of up to trimer-trimer receptor-ligand configurations. Features of this 1^st^ model comprise (i) a direct signal transduction from the extracellular to the intracellular side without the need for massive spatial molecular rearrangements as well as (ii) its occurrence already at low molecular expression levels. A second model proposes TNFRs to form inactive dimers prior to their activation, which in turn assemble into a supramolecular honeycomb lattice^9^. After TNF ligand binding and receptor activation the intracellular receptor domain is cross-linked to reestablish the honeycomb lattice on the intracellular membrane side. Features of this 2^nd^ model are (i) a unique molecular complex permitting robust signal initiation and (ii) potential signal amplification by a factor of ∼ 1.4^10^.

Here, we scrutinize the two models, choosing CD95 as an example of TNFRs, as its exclusive activation by CD95L facilitates data quantification and interpretation. Moreover, qualitative observations of CD95 oligomerization in the cell plasma membrane have been reported^11^, albeit a quantification of oligomer sizes in live cells is missing. This is most likely due to a lack of suitable techniques to discern different oligomerization states during the signaling process. To address this need, we here introduce a strategy based on complementary state-of-the art microscopy and spectroscopy^12,13^ techniques and their further developments as multiscale approach to cover a very large ranges in concentration, time, and space (Figure 1c). In particular, we advance and synergize the readouts of Cell Lifetime FRET Image Spectroscopy (CELFIS), Stimulated Emission Depletion (STED), polarization-resolved confocal Photobleaching Step Analysis (cPBSA), and use Fluorescence Correlation Spectroscopy (FCS). Our strategy also comprises a small library of CD95 variants with different signal initiation competency as well as high-fidelity monomer and dimer controls. In all cases, rigorous image analysis and benchmarking against control samples allowed us to identify concentration or photophysical effects and to quantify CD95 oligomeric states. Thus, we map the regulation of CD95 before and during the whole signaling process and derive a minimal model of CD95 signal initiation. Notably, the presented multiscale toolkit can also be applied to study the oligomerization of other membrane receptor systems, quantitatively.

## Results

### Engineered plasma membrane receptors for molecular quantification in Super-resolution and Multiparametric Fluorescence Microscopy

We have collected a small library of mEGFP and mCherry labeled CD95 variants with different competency to recognize and transduce the signal initiated by CD95L (Figure 2a). Next to monocistronic plasmids, we used bicistronic constructs, combining mCherry and mEGFP labeled proteins, to ensure homogeneous co-expression of donor and acceptor fluorophores during FRET measurements. To quantify receptor oligomerization states, we established high-fidelity monomer and dimer controls using mEGFP or mCherry labeled CD86 and CTLA4 membrane receptors, respectively. As described below, generating a pseudo-dimer control from CD86 with two genetically fused mEGFP was necessary to determine the CTLA4 dimerization state. Further details on the design of the 13 plasmids are found in the Methods section. Prior to measurements, correct integration of all receptors into the plasma membrane was verified using confocal microscopy (see Supplementary Figure 1).

**Figure 2:**
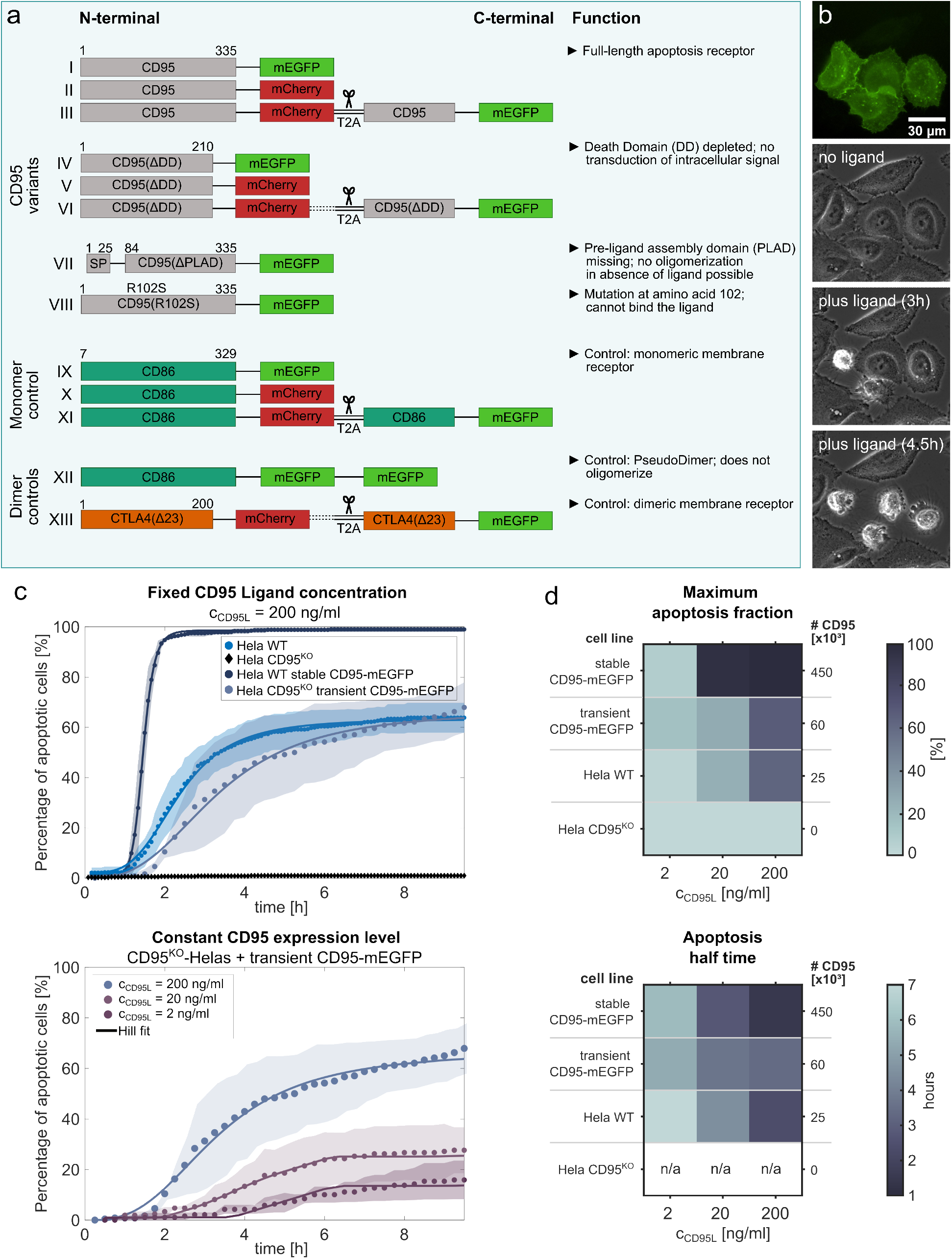
Engineered receptor variants for molecular quantification and characterization of molecular concentration dependent apoptosis dynamics. **a)** Schematic illustration of engineered CD95 variants of different signaling competency (I-VIII) as well as monomer (IX-XI) and dimer controls (XII-XIII). Bicistronic plasmids are used in CELFIS, mEGFP labeled monocistronic plasmids with all other techniques. Numbers refer to the amino acid sequence. Dashed lines indicate optional linkers. Blue panels illustrate the method. **b)** From top to bottom: mEGFP fluorescence and phase contrast microscopy of Hela CD95^KO^ cells transiently transfected with CD95-mEGFP before and after CD95 Ligand addition. 3 h and 4.5 h after 200 ng/ml CD95 Ligand addition apoptosis of transfected cells is observed. Non-transfected cells are unaffected by CD95 Ligand. **c)** Percentage of apoptotic cells over time after CD95 Ligand incubation. From a Hill equation fit (solid line, Equation (1) in Methods) characteristic apoptosis dynamics parameters shown in d) are derived. Top: comparison of cell lines with different CD95 expression level exposed to 200 ng/ml ligand concentration. Bottom: comparison of Hela CD95^KO^ transient CD95-mEGFP cell line exposed to ligand concentrations of c_CD95L_ = 2, 20, 200 ng/ml. Data points show the weighted mean, shaded area the standard deviation of three independent measurements. N >180 cells per sample. **d)** Hill fit parameters: maximum apoptosis fraction (top) and apoptosis half time (bottom) of different cell lines and ligand concentrations c_CD95L_. n/a indicates data where no Hill fit was possible due to a low percentage of apoptotic cells. For further details see text and Methods.

### The efficiency of signal initiation relies on receptor expression levels and ligand concentrations

We first examined CD95 signal initiation and transduction on the cellular level to quantify effects of different ligand concentrations and receptor densities on the signaling kinetics and outcome. To this end, we recorded HeLa cell lines exhibiting different CD95 receptor expression levels of 0 to 4.5 · 10^5^ receptors per cell, as quantified by flow cytometry. Cells were exposed to various ligand concentrations and the kinetics of the cellular fate decision was monitored. Several hours after CD95L incubation, the cells showed typical apoptosis characteristics such as initial blebbing followed by cell shrinkage (Figure 2b). In all cases, the kinetics of apoptosis signaling followed a sigmoidal progression. The initial onset just one hour after ligand addition indicated the minimal time the signal takes from apoptosis initiation until the eventual death of the cell. The predominant time interval of apoptosis events was between 1 to 5 hours after ligand addition, whereas the slowest signaling outcome was detected after 5 to 7 hours, depending on the experimental situation. The few apoptosis events recorded after this time were attributed to naturally occurring apoptosis. We observed a ligand dependent efficiency of apoptosis induction from 3% to 99% apoptotic cells, when the ligand concentration was increased from 2 to 200 ng/ml. Similarly, apoptosis initiation scaled with the number of receptors expressed on the cell surface, where a complete knockout of CD95 (0 receptors) led to no apoptosis, 2.5 · 10^4^ CD95 molecules/cell led to 60-75% apoptotic cells and 4.5 · 10^5^ CD95 molecules/cell led to 99% apoptosis (Figure 2c/d). A fit of the Hill function (see Methods) yielded the time after which half of all apoptotic cells had died. These half-times ranged from 1.5 h to 8 h and became shorter with higher CD95 ligand concentration or receptor cell surface expression (Figure 2d). Cells expressing CD95(ΔDD) and CD95(R102S) served as a negative control and showed apoptotic cells of less than 15% within 10 hours caused by natural apoptosis or potentially transfection stress (Supplementary Figure 2).

For CD95(ΔPLAD), the apoptosis dynamics slightly exceeded the negative controls with up to 25% of apoptotic cells (Supplementary Figure 2). Analyzing the apoptosis kinetics allowed us to define characteristic time points of the signaling process important for subsequent measurements with CELFIS, cPBSA, FCS or STED: (i) time points before signal initiation, (ii) directly after ligand addition, (iii) when most cells underwent apoptosis, and (iv) when all signaling events finished. Moreover, in all apoptosis experiments, the kinetics exhibited a strong correlation with ligand and receptor concentration, demonstrating that signal initiation is highly dependent on the absolute number of activated receptors. For this reason, we payed particular attention to the number of ligands and receptors in the system during the following measurements.

### Ligand induced signal initiation does not affect receptor mobility in the plasma membrane as revealed by live-cell FCS

Prior to single-molecule analyses of CD95 oligomeric states, we tested if CD95 is sufficiently mobile and hence able to form (higher) oligomers using FCS (Supplementary Figures 3&4). Since FCS measurements are more sophisticated in live-cells, due to the natural variability and signal contributions of cytoplasm and plasma membranes, we elaborated an optimized laser power, pinhole and recording time to optimally balance signal-to-noise gains with recordings of less stable fluorophores, such as mEGFP (see Methods and Supplementary Notes 1&2). We recovered diffusion coefficients of CD95 and CD95(ΔDD) in membranes. The obtained diffusion coefficients *D* = 0.23 ± 0.02 µm²/s are typical for individually diffusing membrane proteins^14,15^ and didn’t change in presence or absence of CD95L. The diffusion constants of CD95 were also comparable to those of our control constructs with single and double transmembrane helices, CD86_D0_ and CTLA4_DA_, respectively (Supplementary Figure 3, Supplementary Table 1). Overall, this data confirmed sustained CD95 mobility without significant changes in *D* during the whole signaling process.

### Small spots of receptors below STED resolution and not large CD95 networks govern the distribution in the plasma membrane

We tested the CD95 membrane distribution for local accumulations or supramolecular cluster formation by STED nanoscopy. To this end, we fixed the transfected Hela cells 2h after ligand addition when the signaling was initiated in most cells. CD95-mEGFP was stained with GFP-nanobody Atto647N and the membrane surface was imaged with STED at 40 nm FWHM resolution. STED images revealed a distribution of CD95 in characteristic spots for which we established a quantitative analysis using time-gating with maximum likelihood estimator-based deconvolution followed by a watershed object segmentation and determination of spot size and brightness (Figure 3a/b; see Methods).

**Figure 3:**
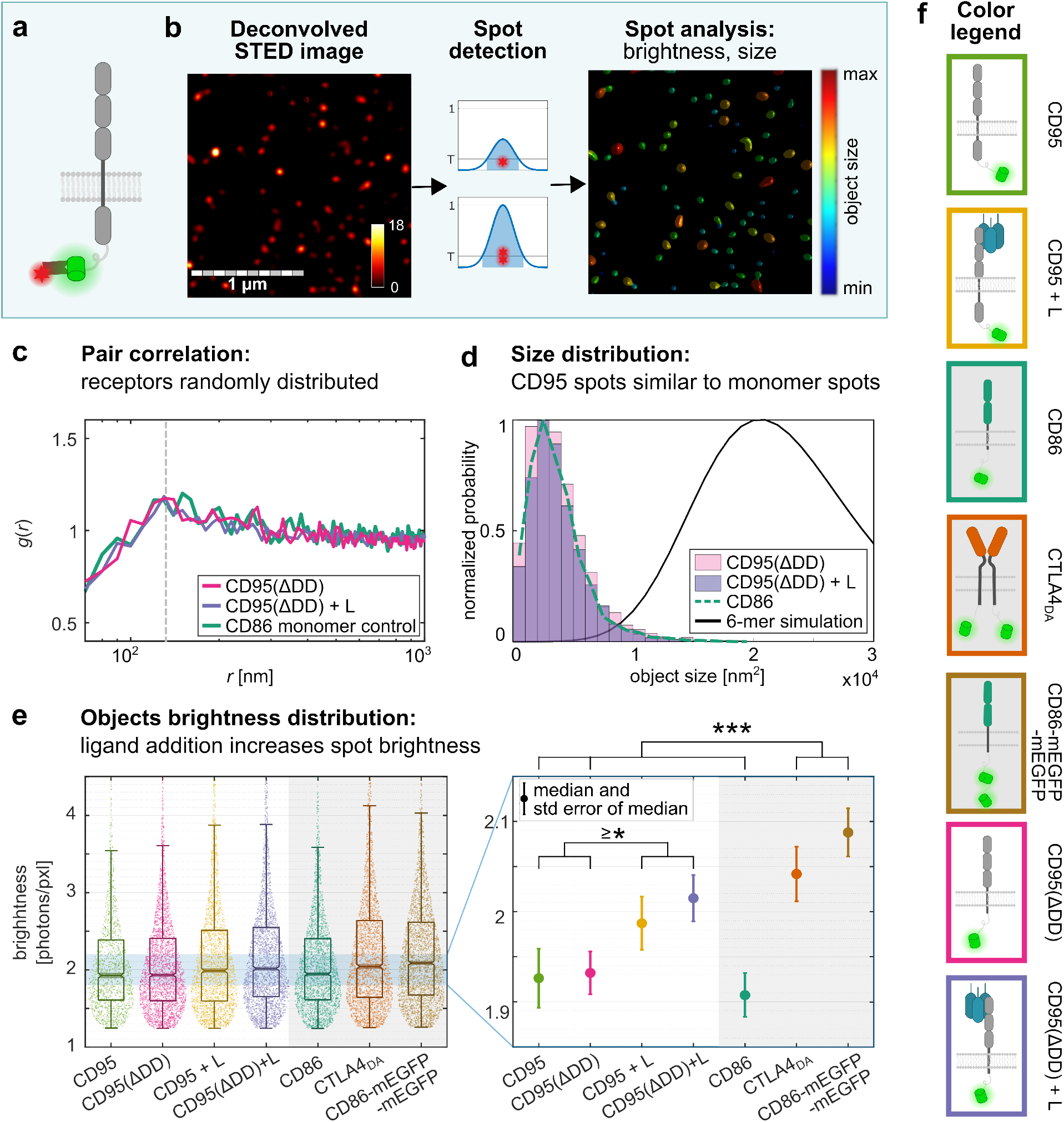
Quantitative STED imaging reveals randomly distributed CD95 spots and systematic changes in object brightness. **a)** Schematic representation of CD95-mEGFP with GFP-nanobody Atto647N labeling. **b)** Exemplary deconvolved STED image (left) of Hela CD95^KO^ transiently transfected with CD95-mEGFP and threshold-based (*T*) spot detection and filtering (middle) followed by spot analysis (right) using Huygens SVI. Blue panels illustrate the method. **c)** Pair correlation function *g(r)* (Equation (3) in Methods) of detected spots for CD95(ΔDD) with/without ligand as well as CD86 monomer control. Distances *r* ≥ 130 nm (right of dashed line) with *g(r)* ≈ 1 indicate a random distribution. The decrease in correlation for *r* < 130 nm arises from finite PSF size effects and no particular distribution (see also Supplementary Figure 5b). **d)** Size distribution of detected spots. CD95(ΔDD) object sizes in the presence and absence of CD95 ligand do not exceed the spot sizes adapted by the CD86 monomer control. Simulation of a 6-mer illustrates the distribution expected for higher order clusters. **e)** Violin -box plots show the distribution of object brightness up to 4.5 photons/pxl for all samples (left). Detail of median brightness (right) reveals a significantly higher value for dimer controls (>2 photons/pxl) compared to CD86, CD95, and CD95(ΔDD) in absence of the ligand. Ligand addition shifts the median brightness of CD95 and CD95(ΔDD) towards the dimer controls. Mann-Whitney U-test with ***: p < 0.001, *: p < 0.05. **f)** Legend: cartoons illustrating the sample receptors. Box color code for each receptor used throughout the manuscript.

To test for higher order pattern within the receptor spot distribution, we first calculated the pair correlation function *g(r)* of the spot centers (Figure 3c). Our data and simulations revealed a random distribution of spots over the membrane surface for all receptors in absence and presence of the ligand. Note, that the decrease in correlation at radii below 130 nm arises from the size of the PSF (for CD95, dimer controls, and simulation see Supplementary Figure 5). From this data, an average concentration of 20 spots/µm^2^ was derived, corresponding to an average distance of 224 nm between spots for an intermediate expression level of about 4 · 10^4^ − 8 · 10^4^ receptors per cell. This estimate is in line with flow cytometry results when only few receptors are assumed for each spot. In addition, the size distribution of CD95(ΔDD) spots before and after ligand addition was comparable to the CD86 monomer control distribution. Simulation of a 6-mer illustrated the distribution expected for higher order clusters (Figure 3d). These data indicated that the existence of higher oligomers/networks was rather unlikely (data of CD95 and dimer controls in Supplementary Figure 5).

To assess this readout further, we evaluated the spot brightness [photons/pixel] of round, resolution-limited spots before and after ligand addition (Figure 3e). Initially, the large spread in spot brightness of CD95 samples was interpreted as the existence of CD95 monomers as well as CD95 oligomers and few higher order networks. However, measurements of monomer and dimer control samples revealed similar distributions. Several reasons could cause spots with varying brightness or sizes exceeding the resolution limit, that question the existence of higher oligomers: (1) local concentration fluctuations, (2) limitations in staining efficiency, (3) photophysical effects, or (4) sample orientation in the membrane. The analysis of the registered polarization-resolved fluorescence revealed a wide range of fluorescence anisotropy values from 0 to 1. This is evidence for a wide distribution of static orientations of stained fixed receptors in the membrane (Supplementary Figure 6). Thus, the observed variations in brightness mainly arise from different orientations of the molecular absorption and emission transition dipole moments.

Under these circumstances, we used the median values of the spot brightness as a robust measure for the distinct samples (Figure 3e). The average monomer brightness was 1.91 photons per pixel (CD86), and the dimer samples were 2.04 (CTLA4_DA_) and 2.08 (CD86-mEGFP-mEGFP) photons per pixel, respectively. The lower brightness of CTLA4_DA_ compared to CD86-mEGFP-mEGFP can be understood from the fact that CTLA4_DA_ dimers consisted of donor-donor as well as of donor-acceptor pairs. A CTLA4 donor only (CTLA4_D0_) expression was not possible, since the plasmid did not localize to the membrane correctly. In addition, and in contrast to the state of knowledge, CTLA4 does not always build 100% dimers, but the dimer fraction depends on the total receptor concentration (as determined by us with CELFIS below). CD95 samples in the absence of a ligand exhibited a median close to the monomer value, whereas after ligand addition, a significant shift toward a median value of 2 was obtained. These results indicate that some oligomers, but no hexagonal networks, consisting of 18 receptors or more, would form. These analyses also highlight the importance of using high-fidelity monomer or dimer controls as molecular benchmarks. In order to determine the CD95 oligomerization state precisely, we then performed cPBSA and CELFIS measurements. These techniques also have the advantage that no additional staining is needed so that an overall higher label density is expected for fluorescent proteins.

### Confocal Photobleaching Step Analysis (cPBSA) identifies sensitive changes in ligand-induced receptor recruitment

Since STED and FCS are not sensitive enough to quantitatively determine the CD95 oligomer fraction and stoichiometry in resolution-limited spots, we also used Photobleaching Step Analysis (PBSA). In the past, PBSA was used to measure *in vitro* samples with photostable organic fluorophore labeling, to determine the number of membrane bound proteins^16^, the degree of Quantum Dot labeling^17^, or the number of fluorescent labels on DNA origami^18^, amongst others. To apply PBSA to CD95, we advanced the technique to be compatible with widely available confocal microscopes and to use it with intracellular fluorescent labels with minimal background noise and without bleaching large areas of the cell (Figure 4a-c). Additionally, the confocal setup gave us full access to spectroscopic tools, which we used here to robustly interpret mEGFP bleaching steps despite the lower photostability and brightness of mEGFP compared to stable organic fluorophores (see Supplementary Note 3, Supplementary Tables 2-4, and Supplementary Figures 7-12). For example, signal fluctuations due to dark states were quantified by computing the cross-correlation function of the polarization-resolved intensity traces, *G*_*ps*_*(t)*, which allowed us to determine characteristic relaxation times for blinking, *t*_b_, and bleaching, *t*_bleach_ (Figure 4d). Although we used circular polarized excitation, polarization effects arising from the presence of static emission dipoles caused variations in the single fluorophore brightness, similar to observations made with STED (Supplementary Figure 10). cPBSA was realized by a fast overview scan of the cell’s lower membrane to identify receptor locations followed by placing a diffraction limited spot at the respective region of interest and recording the bleaching trace (Figure 4a-c; compare Methods). As the fluorophore brightness shifted slightly from day to day due to laser power changes, we calibrated the effect of brightness variations by changing the time bin sizes *in silico*, which is analogous to changing the laser power. This showed that the number of steps scaled with laser power (Figure 4e). Subsequently, we correct for this effect by adjusting the minimal step size by the same factor (compare Methods, Supplementary Table 2 and Supplementary Figure 8). Thereafter, the Kalafut-Visscher (KV) algorithm^18,19^ was used to derive the number of fluorophores per measurement spot.

**Figure 4:**
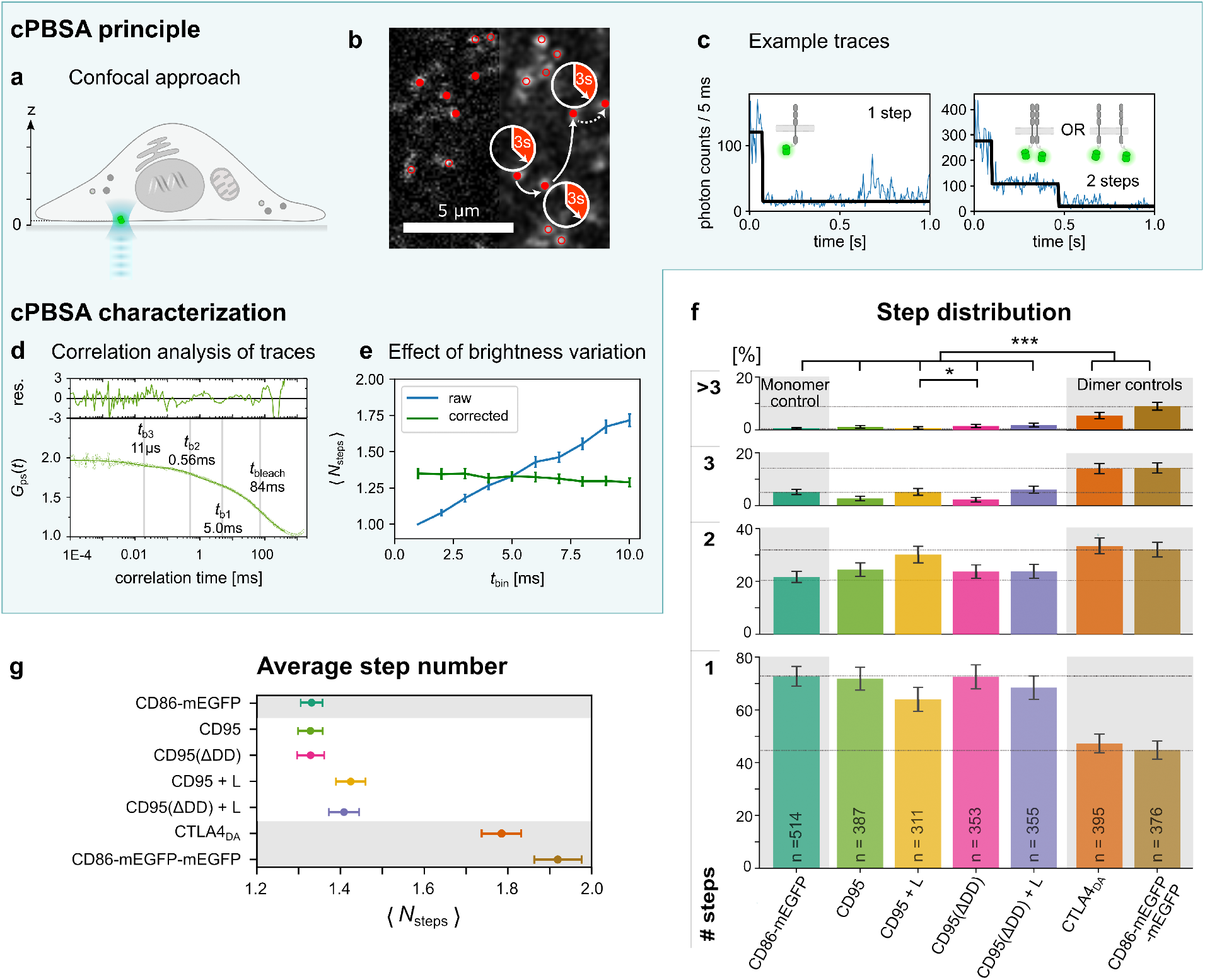
Confocal PBSA reveals the stoichiometry of CD95 in fluorescent spots. cPBSA principle. **a)** The confocal approach enables local trace analysis with minimal overall sample bleaching. Traces may be recorded subsequently at different positions on or inside the cell. **b)** Confocal PBSA spot detection algorithm: acquired confocal overview image (left-half) is smoothed using a Gaussian filter with 1 pixel sigma (right-half). Fluorescent spots exhibiting maxima higher than 4 photons (red circles) and diffraction limited areas with no adjacent neighbors are selected (red dots). Bleaching traces are recorded from each red dot for 3 seconds. **c)** Left: exemplary trace of a monomer. Right: exemplary trace for either a dimer or two monomers in one confocal spot (crowding). **cPBSA characterization: d)** Cross-correlation function *G*_*ps*_*(t)* (Equation (8) in Methods) of CD95 bleaching traces yield characteristic correlation times of mEGFP blinking and bleaching events in cells. Characteristic time scales are derived from a 4 term global fit (see Supplementary Figure 10). **e)** Increasing the time bins (which is analogous to an increased brightness) increases the number of steps found. This is corrected for by changing the minimal step size in the same proportion (see Supplementary Figure 8). **cPBSA results f)** Bar diagram of step occurrence. 1,2 and 3 photobleaching steps were primarily detected. The monomer and dimer controls are used to characterize the fraction of multimer events attributed to crowding. Errors bars are calculated from Poisson statistics. **g)** Mean number of fluorophores and standard error of the mean for data shown in f). A small increase in fluorophore number is detected for CD95 (* with p = 0.026) and CD95(ΔDD) (n.s. with p = 0.169, Mann-Whitney U-test with ***: p < 0.001, *: p < 0.05) in presence of the ligand. The average fluorophore number is significantly smaller compared to those of dimer controls.

In all cPBSA measurements mostly single, double or triple bleaching steps were detected. Only dimer controls exhibited bleaching traces with a higher number of fluorophores per spot. In case of CD95 and CD95(ΔDD) more than 70% of traces exhibited a single step, 23% two steps and about 2% three or more bleaching steps. Upon ligand addition, the fraction of monomers decreased to about 60%, whereas traces of two or three bleaching steps rose to 25% and 5%, respectively (Figure 4f). In absence of the ligand, CD95 and CD95(ΔDD) exhibited a similar distribution of detected fluorophore number compared to CD86 and also the average fluorophore number (⟨***N***_steps_⟩ of 1.33 was identical for these cases. Hence, we concluded, that CD95 is monomeric in its inactive state.

Note, that an elevated average fluorophore number of ⟨***N***_steps_⟩ = 1.33 instead of 1 was found, since also multi-step events corresponding to multiple fluorophores in a single confocal detection volume were recorded (Figure 4g). Intriguingly, these were found in all datasets, including the monomer control dataset. As in case of our STED data, such events may arise from true oligomerization as well as molecular accumulation due to local concentration fluctuations. To estimate effects of molecular proximity within the confocal volume on the appearance of multi-step traces, we calculated an occupancy probability based on the signal density above a particular threshold (Supplementary Figure 8). For this, a weak correlation with ⟨***N***_steps_⟩ was found, supporting the concentration fluctuation hypothesis. We further verified, that the occupancy probability distribution was comparable between samples, such that no additional correction of traces had to be introduced. After ligand incubation, a slight shift to higher oligomerization states was observed for CD95 (+7%) and CD95(ΔDD) (+6%) with an average fluorophore number rising to 1.42 (Figure 4f/g). To interpret this change in light of the appearance of local concentration fluctuations or photophysical effects, we rated it against the dimer controls CTLA4_DA_ and CD86-mEGFP-mEGFP. The two-step controls were significantly higher than all other measurements (p < 0.001) with CD86-mEGFP-mEGFP and CTLA4_DA_ exhibiting ⟨*N*steps⟩ of 1.92 and 1.78, respectively. The value for CTLA4_DA_ was slightly lower than for CD86-mEGFP-mEGFP for the same reasons mentioned in case of STED. Both values were also smaller than the expected value of 2, most probably due to the maturation efficiency for mEGFP being ≲ 80% ^16,20^, but, on the other hand, the difference to the monomer control is higher than for nanobody staining because no additional preparation step is needed. Overall, cPBSA analyses show that few CD95 receptors accumulating in spots are sufficient to trigger apoptosis effectively.

### CELFIS reveals for a large receptor concentration range that 8-17% monomers becoming part of dimers and trimers suffice for efficient apoptosis induction

Finally, since the above techniques are not capable to distinguish molecular proximity from intermolecular interactions within diffraction-or STED resolution limited spots and have limited capacity to probe variability in biological phenotype, we used and advanced FRET to probe transfected cells with different receptor surface concentrations.

As before, CD95 and CD95(ΔDD) were measured in absence and presence of the ligand. The monomeric receptor CD86 served as a negative no-FRET control and CTLA4 as a dimeric positive control. In all cases, bicistronic plasmids were used to ensure homogeneous donor and acceptor expression. Figure 5a shows the localization of the CD95 receptor in live cells by confocal images on the lower cell membrane. The increased intensity at cell edges and cell-to-cell contacts confirms the primary integration of the receptor into the cell plasma membrane. Similar images were recorded for CD95(ΔDD), CD86 and CTLA4 (see Supplementary Figure 1).

**Figure 5:**
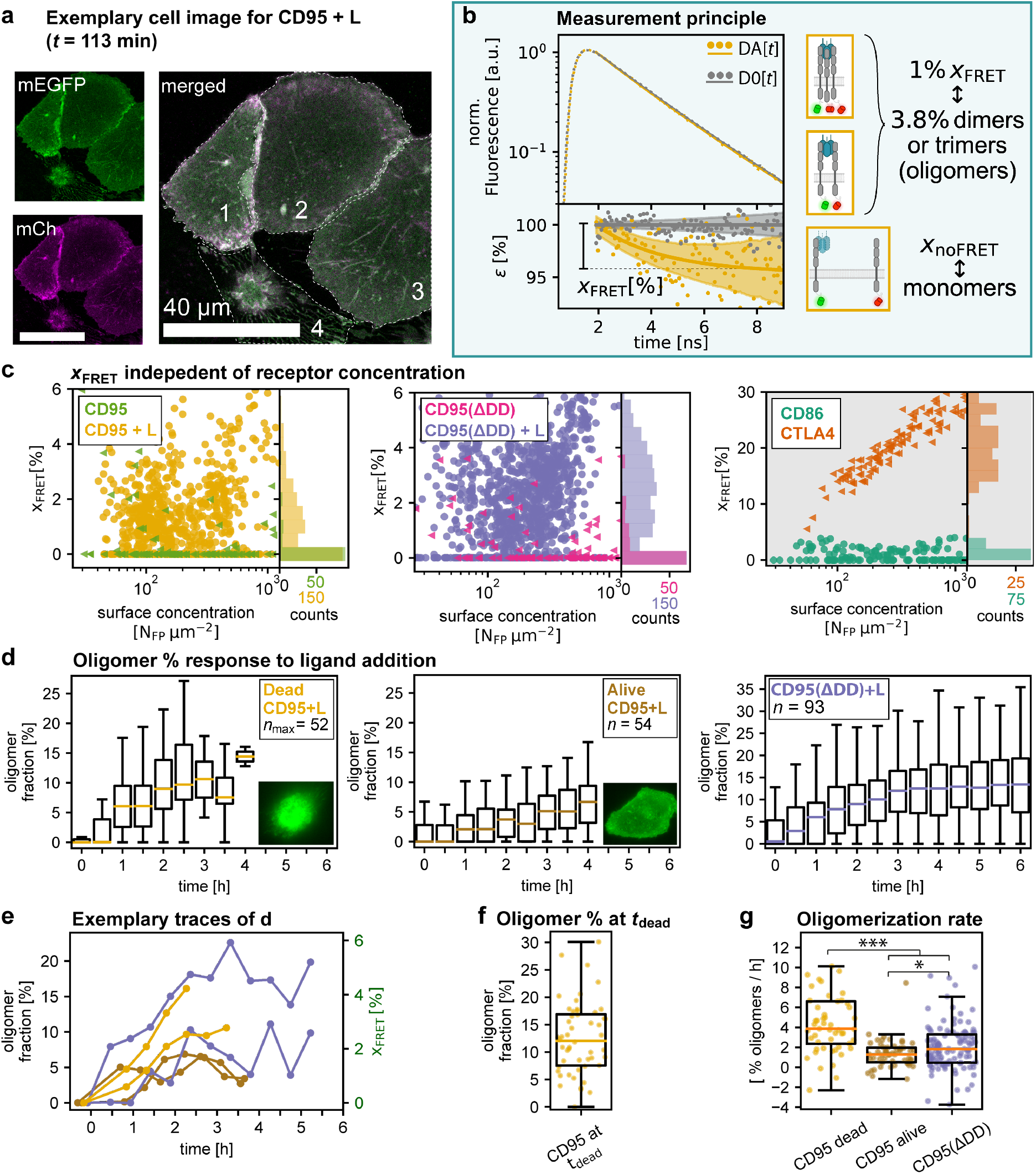
CELFIS quantifies the CD95 oligomerization state over a large concentration range. **a)** Confocal fluorescence image indicating correct integration and colocalization of mEGFP and mCherry labeled CD95 in the membrane. Cells 1, 2 and 3 are alive at the time of measurement, whereas cell 4 already underwent apoptosis. Fluorescence lifetimes were recorded over the whole cell membrane for each cell. **b) Measurement principle:** Top: fluorescence lifetime distribution for a live cell in absence (D0) and presence (DA) of FRET. Bottom: normalized fluorescence decay ε shows the quenched fluorescence fraction (x_FRET_) in presence of the acceptor due to FRET. Conversion of x_FRET_ into oligomer fraction was realized by a theoretical x_FRET_ determination of a pure dimer sample (see text). Blue panels illustrate the method. **c)** x_FRET_ histogram and scatter plot as a function of receptor surface density. CD95, CD95(ΔDD) and the monomer control CD86 are monomeric over the whole concentration range. After CD95 ligand incubation, a small fraction of CD95 and CD95(ΔDD) oligomerizes to dimers or trimers. Intriguingly, CTLA4 switches from a monomer to a dimer with increasing receptor concentration. N > 324 cells from at least 4 independent experiments were analyzed per condition. **d)** Dynamics of oligomer formation after CD95 ligand addition. The oligomer fraction was calculated from repeated measurement of the same cells and averaging over many traces. Boxplots are shown with colored medians. Oligomer fractions saturate after 3-4h. **e)** Exemplary evolution of the oligomer fraction in single cells over time. Legend same as in d). **f)** Boxplot of the oligomer fraction at the last time point before apoptosis. **g)** Oligomerization rate over the first 3 hours or less due to the timepoint of apoptosis. Legend same as in d). Mann-Whitney U-test with ***: p < 0.001, *: p < 0.05.

To systematically tune the range of receptor surface concentrations and to thereby obtain insights about the molecular concentration fluctuations suggested by the above techniques, we titrated the amount of receptor DNA used for transfection against an empty vector, while keeping the total amount of DNA constant. We further determined the molecular brightness of the fluorophore to convert fluorescence intensities into surface densities, ***N***_FP_/µm². Here, changes of the donor fluorophore lifetime due to FRET only occurred, if receptors labeled with a mEGFP donor and a second receptor with a mCherry acceptor molecule were in close proximity due to binding (< 10 nm).

For Cell Lifetime FRET Image Spectroscopy (CELFIS), we evaluated the data of receptors on the lower cell membrane and integrated all photons over the cell bottom surface in a single fluorescence decay per cell to determine the average oligomerization state with great accuracy. Figure 5b illustrates the core principle of CELFIS: the fluorescence decay was measured in the FRET sample (DA) as well as the control sample, expressing the donor in absence of the acceptor (D0). Normalization of the DA fluorescence decay with respect to the average D0 decay allows one to extract the FRET-induced donor decay (*ε*_D_*(t)*), equations (9-12)^21-23^. Its amplitude drop directly corresponds to the donor fraction, *x*_FRET_, that was quenched by FRET^22^.

We determined *x*_FRET_ values for each cell individually and studied its dependence on the receptor surface concentration ***N***_FP_/µm² (Figure 5c). Thereafter, we benchmarked the data of CD95 against signals obtained from the CD86 and CTLA4 controls. As expected, we observed no FRET for CD86 which was predominantly monomeric up to a concentration of 1250 receptors/µm². At this point, a systematic increase in FRET indicates the onset of proximity FRET, which was also observed for CD95 and CD95(ΔDD) in absence of the ligand. For this reason, and since proximity FRET was suggested to lie in this concentration range^24^, we evaluated the FRET data only up to the threshold of 1250 receptors/µm². FRET measurements of CD95 and CD95(ΔDD) without ligand likewise showed that both receptors are monomeric. Upon ligand addition, the value of *x*_FRET_ increased immediately by a few percent. Together with our cPBSA results, these values suggested formation of dimers and/or trimers (Figure 5c). Finally, we derived a relation to approximate the oligomer fraction from the measured *x*_FRET_ by calculating a sample-specific maximum FRET signal *x*_FRET,max_ for a purely dimeric sample (see Methods). This calibration accounts for i) the distance distribution between the two fluorescent proteins with long linkers^21,25^ (see linker list in Supplementary Table 5), ii) the abundance of no-FRET species due to donor-donor dimers and iii) an estimated maturation efficiency of 80% for EGFP and mCherry^16,20^, yielding a *x*_FRET,max_ of 29% and 26% for CTLA4 and CD95, respectively. Hence, for the CD95 protein, a pure dimer sample (100% dimers) corresponded to 26% *x*_FRET_ and, equally, 1% *x*_FRET_ corresponded to a ∼ 3.8% oligomer fraction. The calculation for CTLA4 was analogous.

Equipped with these tools, we then probed how the oligomerization state changed over time until the point of apoptosis. Here, we recorded FRET data over 0 to 6 hours after ligand addition by repeated measurements of the same cells. Cells expressing the full-length CD95 were classified according to whether apoptosis occurred within the observation time of 4h (Figure 5d). For those that underwent apoptosis, the oligomer fraction started close-to-zero and increased quickly up to an 8% median value, whereas cells that did not show apoptosis exhibited a slower oligomer formation, reaching a ∼ 5% median after 4h. CD95(ΔDD) expressing cells, where downstream signaling was suppressed, showed a slightly higher initial oligomer fraction and reached a population equilibrium of 12% median after ∼ 3h. In individual cell traces rising and/or falling oligomer fractions were detected (Figure 5e), representing transient CD95 dimerization or binding/unbinding kinetics of CD95 to CD95L (see Methods for further analyses). As a measure of CD95 oligomerization needed to initiate apoptosis, the oligomerization fraction just prior to apoptotic blebbing and shrinkage was estimated, amounting to the interquartile range of ∼ 8 to 17% with a median value of 12% (Figure 5f). Finally, we determined the oligomerization rate from the oligomer fraction change per time interval, which was faster in case of CD95 transfected cells that died (3.9% oligomers/h) compared to CD95 or CD95(ΔDD) transfected cells which stayed alive (with 1.3% and 1.8% oligomers/h respectively, Figure 5g). We further investigated the oligomeric state in membrane areas classified according to their brightness, revealing that the oligomerization is not limited to certain areas and occurs according to its concentration dependence (see Supplementary Note 5 and Methods). Overall, our results demonstrate that oligomers form within 2 - 3 hours over the whole membrane. Oligomerization requires ligand addition and can develop in absence of a death domain, indicating that CD95 oligomerization may be mediated by the transmembrane domain only in the receptor activated state, as previously suggested^26^, or simply via ligand binding. Finally, only about ∼ 8 - 17% oligomers in the form of dimers or trimers are necessary for efficient signal initiation.

## Discussion

Here, we present an advanced molecular sensitive imaging toolkit combined with multiscale analysis to decipher the spatiotemporal organization and dynamical interactions of CD95 during signaling in the cell plasma membrane. We determine CD95 oligomerization states and find receptors to be initially monomeric and homogeneously distributed on the cell plasma membrane. In previous studies TNFRs (including CD95) were reported to appear as monomers, dimers or trimers in the absence of a stimulus^8,27^. Pre-ligand dimer- and mostly trimerization of CD95 was reported in several works, where receptors were purified and reconcentrated (e.g. ∼ 0.5 mg protein/ml^28^) from E.coli or mammalian cells and analyzed by gel filtration, western blot or crystallography^8,28-30^. In three further studies based on crystallography and NMR spectroscopy, CD95 was suggested to form higher oligomeric structures of penta-or hexagonal shape in bicelles (Figure 6a). In contrast to these biochemical *in vitro* approaches that can affect structural features in membrane proteins^31^, molecular sensitive imaging of receptors directly in the cell plasma membrane revealed primarily monomer and dimer formation^27,32^. Our data obtained in live cells without fixation and staining confirms the latter results and suggests that the situation in the native membrane environment, with small or no oligomers developing, is significantly different from the purified receptor case (Figure 6).

**Figure 6:**
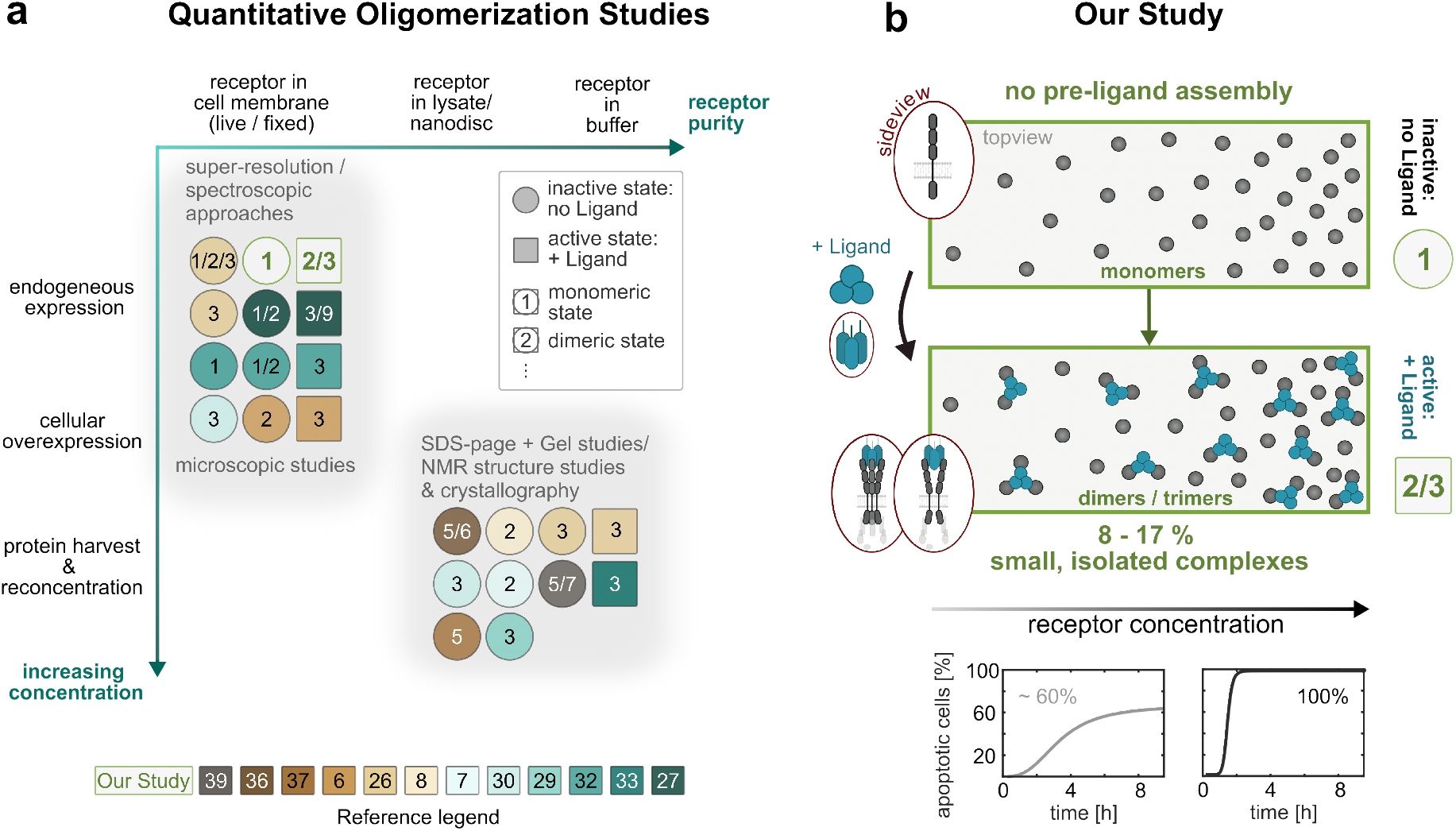
CD95 oligomerization state over a large concentration range. **a)** Summary of studies on quantitative TNFR oligomerization in context of the measurement parameters receptor concentration and receptor environment for different measurement technique. Numbers in circles or boxes indicate the measured oligomerization grade. Numbers in the reference legend correspond to publications in the reference list. **b)** Schematic illustration of the minimal model of CD95 signal initiation shown in this study: monomeric receptors (no pre-ligand CD95 assembly). After ligand binding 8 to 17% of receptors form small, isolated complexes. Increasing receptor concentrations (surface expression level) do not lead to higher oligomer fractions. Since a higher number of apoptotic cells is obtained with increasing receptor concentration, the absolute number of active, oligomerized CD95 appears as a decisive parameter.

After ligand addition, we find dimers and trimers forming within the first 2-3 hours with a final fraction of 8 to17% receptors exhibiting oligomerization. Interestingly, the majority of previous studies reports CD95 and other TNFRs to be trimeric after ligand addition. Among these, molecular sensitive techniques, such as crystallography, single molecule localization microscopy, and biochemical receptor cross-linking studies favor the trimeric state^3,6,7,27,33^. A general observation of molecular clustering was also reported in widefield fluorescence microscopy studies, albeit without quantifying molecular numbers or interactions^27,34,35^. Now, being equipped with the toolkit to quantify molecular oligomerization as presented herein, it would be interesting to also reconsider these cases.

From a structural point of view, three types of molecular interactions are currently discussed to give rise to TNFR signaling and to explain the reported observations: (i) the direct coupling of up to three receptors to the ligand, without the need of direct intermolecular interactions between receptors, (ii) interactions between CD95 transmembrane domains after ligand activation^26^, and (iii) intracellular crosslinking of two CD95 DDs via FADD^36,37^. Cases (i) and (ii) would result in close packing of CD95 receptors with few nm intermolecular spacing around the ligand up to a trimer-trimer configuration^38^. Case (iii) suggests that recruitment of FADD and interaction with the DD results in crosslinking of two DDs. If a crosslinking between different trimer-trimer units occurs, also the higher oligomeric structure of hexagons could develop, placing the receptors some ∼ 12 nm apart (with exact values varying between TNFRs)^9,10,36,39^. Yet, the DD-FADD interaction was reported to be weak^36^ and may not occur at low CD95 and FADD concentrations. This may explain the appearance of higher oligomeric structures when purified and reconcentrated CD95 and FADD were investigated^39^. Moreover, as shown in our study, full length CD95 exhibited near identical oligomerization behavior compared to DD truncated receptors, demonstrating that efficient signaling is possible in the absence of DD-DD crosslinking (Figure 5c,d). This leads to the conclusion that the observed CD95 dimer/trimer formation is mediated via direct ligand (i) or ligand-induced transmembrane (ii) interactions.

The difference in oligomeric states found in case of purified receptors relative to cell membrane samples underscore the importance of the physical and molecular environment in which CD95 is measured. This is not surprising, as already molecular mobility and consequently any interaction probability is highly different in purified samples compared to CD95 embedded in the cell plasma membrane (e.g. protein membrane diffusion of D ∼ 0.2 µm^2^/s versus protein diffusion in solution D ∼ 50 µm^2^/s^40^). More importantly, molecular concentration and environment will influence the oligomerization state. In case of the purified samples in presence of detergents a rather high sample concentration of ∼ 100 µM was reported ^36^. In cell lines, we determined molecular expressions to 10 to 1000 receptors / µm^2^, where the lower limit marks the physiological expression level and the upper limit concentration regime is already found in *in vitro* studies. Despite this broad range of concentrations covered in live cells, our data did not show signatures of higher oligomers, suggesting that either concentrations are still too low or that CD95 in contrast to other TNFRs does not form any hexagonal network. Indeed, previous *in vitro* studies of purified TRAIL coupling to Death Receptors 4 and 5 reported changes of molecular stoichiometries in the protein complex only upon increasing molecular concentrations by orders of magnitude from 1 nM to 10 µM^41^. Hence, we conclude that higher oligomerization states of CD95 without ligand may only develop at very elevated receptor concentrations or under conditions, where the hydrophobic region of the receptor such as the transmembrane helix is not fully immersed in a lipid membrane layer, e.g. bicelles^26^ or in a particular cell membrane environment^41^.

While no significant changes in molecular oligomerization are detected, there is a remarkable change in signaling dynamics and the percentage of apoptosis events depending on the absolute ligand and receptor number. Here, as well as in previous studies^34,42^, using different cell types and CD95 expression levels between 5 · 10^10^ − 450 · 10^10^ receptors/cell, a significant acceleration of downstream signaling and systematic increase of apoptosis events was shown when receptor or ligand concentrations were increased. Hence, tuning the absolute number of activated receptors turns out to be a crucial aspect in apoptosis signal initiation.

To provide the above insights, we assembled a multiscale toolkit to cover spatial, stoichiometric and temporal resolution needed for studying receptor oligomerization. The toolkit consists of six techniques including super-resolution and multiparametric fluorescence imaging which were advanced to record data with single-molecule sensitivity. In particular, we established a quantitative spot analysis of STED data, verified receptor mobility with FCS, and determined CD95 stoichiometries in fluorescent spots from cPBSA. In case of the latter, mEGFP fluorescence labeling as well as confocal instead of Total Internal Reflection Fluorescence imaging was established, making cPBSA measurements applicable to common biological samples and more flexible in space, respectively. The automated workflow for time-resolved FRET image spectroscopy in live cells (CELFIS) was developed by the authors during the course of this study to measure and analyze large numbers of cells to obtain the required precision and sensitivity to determine oligomerization states over the whole cell and during the signaling process. Our study highlights the need for parallelized measurements using complementary techniques (in terms of their spatio-temporal resolution and molecular concentration detection) to probe a high dynamic range (µs to hours, nm to 100 µm scales, 1 to 10^@^ molecules/µm^6^) Finally, benchmarking CD95 data against robust monomer and dimer controls, revealed that intense regions on the membrane initially associated with higher oligomerization states may simply arise from molecular concentration fluctuations across the membrane.

To our best knowledge, this study is the first to report a minimal model of CD95 signal initiation identifying 8 - 17% CD95 monomers oligomerizing to dimers and trimers as efficient apoptosis signal inducers in live cells (Figure 6b). Our results do not exclude the existence of proposed higher order oligomeric states, but confirm that they are not necessary in the studied cellular context. In this respect, our study highlights the importance of molecular concentration level determination as well as the use of high-fidelity monomer and dimer controls for quantitative molecular imaging. Our study not only elucidates the debate about CD95 signal initiation mechanisms but also reports strategies of single molecule quantification in live cells, which are generally important for the study of cell signaling processes.

## Methods

### Sample preparation

#### Plasmids, molecular cloning and stable cell lines

For all measurements with transient transfections, a stable Hela cell line with knockout for CD95 was used (HeLa CD95^KO^). It was generated using CRISPR/Cas9^43^, the guide RNA was CATCTGGACCCTCCTACCTC^32^. For apoptosis dynamics, we additionally used Hela WT cells (purchased from the American Type Culture Collection (ATCC, Manassas, Virginia, USA)) and a stable, overexpressing cell line HeLa CD95-mEGFP, expressing CD95-mEGFP on top of endogenous CD95^32^. Hela CD95^KO^ and Hela stable CD95-mEGFP cell lines were kindly provided from *Joël Beaudouin* (formerly IBS, Grenoble).

For CD95 constructs, four different sequences were used: the full-length protein CD95 (amino acids 1-335), a death domain truncated version CD95(ΔDD), CD95(R102S) and CD95(ΔPLAD). For CD95(ΔDD) amino acids 211-335 were truncated. CD95(ΔDD) is not capable to transduce the intracellular signal and is hence ideally suited for long-time observations after ligand incubation as well as to probe oligomerization mediated by the extracellular and transmembrane domain of CD95. CD95(ΔPLAD) is the PLAD (pre-ligand assembly domain) depleted variant, missing amino acids 26-83. It may be used to detect pre-oligomerization based on transmembrane and intracellular interactions. All amino acid numbers refer to the premature protein sequence (including signaling peptide). CD95(R102S) exhibits a mutation at amino acid 102 (pre-mature protein) and is suitable as control that cannot bind the ligand.

As monomer control plasmid, the full-length sequence of CD86^44^ was used. For the dimer control CTLA-4, the last 23 amino acids of the sequence were removed in order to reduce the internalization of the receptor and to concentrate it at the plasma membrane^45^. As a second (pseudo-) dimer control, CD86 was fused to two consecutive mEGFPs. The UniProtKBs of CD95, CTLA4 and CD86 are P25445, P16410 and P42081-3, respectively.

All plasmids except CD86-mEGFP-mEGFP were as well kindly provided from *Joël Beaudouin* (formerly IBS, Grenoble). These plasmids were designed by fusing the coding sequences of the respective proteins C-terminal (intracellularly) via a linker to mEGFP (called D0 / Donor only) or mCherry in the pIRESpuro2 vector (Clontech)^32^ (for more linker details see Supplementary Table 5). Besides these monocistronic constructs for CD86, CD95, CD95(ΔDD) and CTLA, we additionally used bicistronic plasmids combining the mCherry and mEGFP versions of a protein into one plasmid for FRET measurements to ensure homogeneous co-expression of donor and acceptor (called DA / donor-acceptor), where mCherry is first transcribed and thus more abundant. The bicistronic constructs with a 2A peptide use the sequence EGRGSLLTCGDVEENPGP as linker between the two proteins^32^. Note, that solely CTLA4_DA_ was used instead of CTLA4_D0_, as the latter did not localize to the membrane exclusively.

The CD86-mEGFP-mEGFP pseudo-dimer control was synthesized using a cloning service (*BioCat GmbH* Heidelberg, Germany) by fusing two linked mEGFP proteins C-terminally to the CD86 full-length sequence of CD86 in a pcDNA3.1(+) vector (*BioCat GmbH*).

#### Cell culture, transfections and ligand incubation

All cells were maintained in culture medium, consisting of DMEM (Dulbecco’s Modified Eagle Medium) + GlutaMAX™ (31966021, Gibco, Life Technologies Inc., Carlsbad, California, USA) containing 10% FBS (fetal bovine serum) (10500064, Gibco) and 1% penicillin/streptomycin (P/S) Solution (P0781, Sigma-Aldrich, Merck KGaA, Darmstadt, Germany), in an environment with 5% CO_2_ (v/v) at 37 °C.

For all live cell measurements as well as cPBSA, cells were trypsinized (T3924, Sigma-Aldrich) and seeded in an 8-well glass bottom slides (#80827, ibidi GmbH, Gräfelfing, Germany) with a density of 3-5 × 10^4^ cells per well. For STED immunostaining, 100-150 × 10^4^ cells were seeded on a sterile glass coverslip (13 mm diameter, No. 1.5H, 0117530, Paul Marienfeld GmbH & Co.KG, Lauda Königshofen, Deutschland).

Transfections were obtained using ViaFect™ Transfection Reagent (#E4981, Promega Corp., Madison, Wisconsin, USA) at a cell density of 60-70% following the manufacturer’s protocol. For Apoptosis Dynamics, FCS, STED and cPBSA the cells were transfected with 25 ng of target DNA and 975 ng empty vector (pIRES-puro2 or pcDNA) for all used plasmids per two wells of an 8-well slide or one coverslip. For FRET measurements, the bicistronic plasmids were transfected using varying amounts of target DNA to cover a broad range of expression levels: for a transfection in 2 wells, the combinations 25 ng target DNA + 975 ng empty vector, 100 ng target DNA + 900 ng empty vector, 250 ng target DNA + 750 ng empty vector as well as 1000 ng target DNA (no empty vector) were used. Donor only controls (the monocistronic mEGFP fusion version of the proteins) were expressed at these varying concentrations as well.

Live experiments or fixations were done 48-72 hours after transfection. For all live-cell experiments (time-lapse imaging, FCS and FRET), the cells were incubated in Leibovitz’s L-15 Medium (21083027, Gibco) without phenol red, supplemented with 10% FBS (10500064, Gibco) and 1% P/S (P0781, Sigma-Aldrich).

For all apoptosis experiments including the CD95 Ligand, the *FasL, soluble (human) (recombinant) set* (ALX-850-014-KI02, Enzo Life Sciences Inc., Loerrach, Germany) was used. The ligand was prepared according to the manufacturers protocol and further diluted in the respective cell culture or imaging medium. The provided enhancer was used for all experiments except FCS. For experiments using the Enhancer, the enhancer concentration was always 100-fold higher than the ligand concentration. For all apoptosis experiments except the apoptosis dynamics, the ligand concentration was 200 ng/ml.

#### CD95 Quantification by Flow Cytometry

The quantitative CD95 expression level of Hela WT, Hela CD95^KO^ and Hela WT stable CD95-mEGFP was assessed using the QIFIKIT® for quantification of cell surface antigens by flow cytometry (K007811-8, Agilent Technologies, Inc., Santa Clara, California, USA) on a MACSQuant Analyzer 10 (Miltenyi Biotec) following the manufacturer’s protocol accurately. For CD95 detection, a monoclonal CD95 antibody (130-108-066, Miltenyi Biotec B.V. & Co. KG, Bergisch Gladbach, Germany) was used. As negative control an antibody against CD28 was used (70-0281, Tonbo™ A Cytek® Brand, San Diego, California, USA). As the secondary FITC antibody provided with the QIFIKIT® interfered with the mEGFP of the stably expressing CD95-mEGFP HeLa cell line, a secondary anti-mouse antibody conjugated to APC (17-4010-82, eBioscience™, Invitrogen) was used for all samples instead. The measurement was repeated two times independently. For Hela CD95^KO^ with transient CD95-mEGFP, the number was not obtained from flow cytometry but from STED imaging spot density.

#### Cell fixation and Immunostaining

For cPBSA and STED immunostaining, cells were fixed after transfection within the respective seeding vessel (see Section *Cell culture and transfections*). For experiments including the CD95 Ligand (Enzo Life Sciences Inc.), the ligand was incubated for 2 hours at 37 °C before the fixation.

Before fixation, cells were washed three times with cold washing buffer (HBSS (14025050, Gibco) containing 0.1 M sucrose (57-50-1, Carl Roth GmbH + Co. KG, Karlsruhe, Germany) and 1% BSA (A1391, ITW Reagents, AppliChem GmbH, Darmstadt, Germany)). The fixation was obtained using 4% methanol-free formaldehyde (28906, Thermo Scientific, Life Technologies Inc.) in washing buffer for 10 minutes, shaking at RT. For STED, the fixation buffer additionally contained 0.1% Glutaraldehyde (25% in H_2_O, G5882, Sigma-Aldrich), which was not used for PBSA in order to reduce the fixation related green autofluorescence of the sample. Afterwards, cells were washed three times again.

For cPBSA, as a last step, the cells were incubated with 750 mM Tris (Tris(hydroxymethyl)aminomethane, 103156X, VWR Chemicals, VWR International GmbH, Darmstadt, Germany) in DPBS (14190144, Gibco) to quench the autofluorescence of the formaldehyde. Afterwards, they were washed with DPBS (covered with DPBS for the experiment.

For STED immunostaining, the next step was permeabilization with the washing buffer including 0.2% Saponin (47036, Sigma-Aldrich) as permeabilizing reagent for 10 minutes. After 2x washing, the sample was blocked using a blocking buffer (HBSS with 0.1 M sucrose and 4% BSA) for one hour. For the staining step, the GFP-Booster Atto647N (gba647n-100, ChromoTek GmbH, Planegg-Martinsried, Germany) was diluted 1:200 in the blocking buffer and again incubated for 1 hour. Next, extensive washing was done using the washing buffer at least 3 times. As a last step, the coverslips were mounted upside down on a microscope slide using *ProLong™ Diamond Antifade Mountant* (P36965, Invitrogen, Life Technologies Inc., Carlsbad, California, USA) and stored over night before imaging.

## Methods

### Time-Lapse imaging for apoptosis dynamics

The time-laps measurements were performed on an IX83 inverted epi-fluorescence microscope system (Olympus Europa SE & CO. KG, Hamburg, Germany) (details in section *Microscope setups*) using either a 20x oil-objective (NA 0,85, UPLSAPO20xO) or a 60x oil-objective (NA 0.65–1.25, UPLFLN60XOIPH) on a temperature-controlled on-stage heating system (PeCon GmbH, Ulm, Germany) at 37 °C. The CD95 Ligand (Enzo Life Sciences Inc.) (Section *Cell culture, transfections and ligand incubation*) was added to the cells to the desired final concentration on the microscope. Time-lapse videos were acquired with the *CellSense Dimensions* Software (Olympus) by sequential imaging of the phase-contrast channel and, if available, the mEGFP channel (excitation 470/40 nm, emission 525/50 nm) at multiple positions every 5 to 15 minutes over 10 hours. Image analysis was performed with Fiji^46^, using an intensity-based threshold to the fluorescence channel in order to detect successfully transfected cells. Apoptotic cells were manually identified via the phase-contrast channel.

For a mathematical description of the sigmoidal apoptosis dynamics curves *P(t)*, they were fitted (MATLAB R2019a, The MathWorks, Inc.) using the hill equation to characterize the dynamics and cooperativity of the cell response:

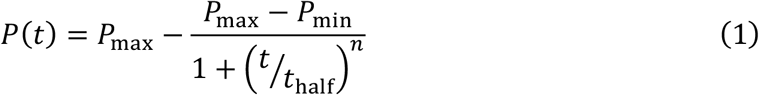

*P*_min_ and *P*_max_ are the minimal and maximal fractions of apoptotic cells and *t*_vpwx_ is the characteristic time after that half of all apoptotic cells died. The hill coefficient (also cooperativity coefficient) *n* indicates the efficiency of the signal induction.

### STED imaging and analysis

STED images were recorded on the Abberior Expert Line Setup (Abberior Instruments GmbH, details in section *Microscope setups*). All immunostained samples (section *Cell fixation and Immunostaining*) were imaged with a 640nm excitation laser (5.3 µW) and a 775 nm STED depletion laser (41 mW) using an oil-immersion objective (NA 1.4, UPLSAPO 100XO, Olympus Europa SE & CO. KG). Before the measurements, channel alignment was performed manually using TetraSpeck Microspheres (T7279, Invitrogen). ROIs of 5 µm x 5 µm (10 nm pixel size, 4.00 µs dwell time, 5 frames) of the bottom cell membrane were recorded.

### Deconvolution & object analysis on STED data

As a first step of data processing, time-gating of the first 2.2 ns was employed to increase the achievable resolution using the home-built programm AnI. The sum of the parallel and perpendicular polarized images was used for further analysis. For deconvolution and image data analysis, Huygens Professional (HuPro Version 21.10.1p2 64b, Scientific Volume Imaging B.V., Hilversum, Netherlands) was used. The deconvolution was performed using the CMLE (Classic Maximum Likelihood Estimation) algorithm with a signal-to noise ratio (SNR) of 3. The convergence stop criterium was set to 0.01 or a maximum of 40 iterations. The automatic background estimation was used with a search area of 0.7 μm radius. After deconvolution, the Object Analyzer of Huygens was used to quantify the object properties of the membrane protein spots. The global object threshold was 1.2 with a seeding level of 1.3, the garbage volume was 2 voxels. Objects touching the image border were excluded from the analysis and only objects with an aspect ratio 0.9 < *D*_x_ /*D*_y_ < 1 .11 of the diameters D in x and y were considered as elongated objects result from crowding. It was verified that this sphericity filter did not preferentially filters large objects.

The size distribution of a 6-mer was simulated by multiple (5x) convolution of the monomer control size distribution with itself.

### Spot Anisotropy Analysis

The spot intensities of the parallel (P) and perpendicular (S) channel, *I*_P_ and *I*_S_, were determined with an individual object analysis of both images (compare Chapter before). The steady state anisotropy *r* was calculated with

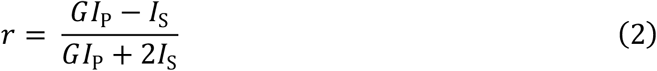

where the polarization correction factor *G* = *η*_P_/*η*_S_ corrects for the instrument’s polarization dependent transmission. *η*_P_ and *η*_S_ are the detection efficiencies of the parallel and perpendicular detection channels. The polarization correction factor *G* was determined to be 0.905.

### Pair correlation

The distribution of object points was analyzed using the pair correlation function *g(r)*^47^:

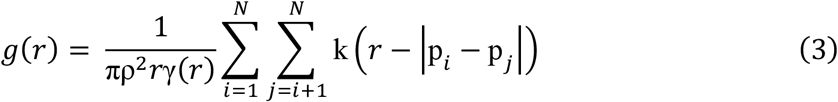

where ρ is the object density in the image, and ÉpÑ − pÖÉ is the distance between two object points p with two-dimensional position *(x, y)*. The object positions were assumed to be planar. The covariance function *γ* and kernel k are defined in^48^.

The pair correlation of the objects found by the Huygens Object Analyzer was calculated using a locally designed MATLAB script (R2019a, The MathWorks, Inc.) following the example of ^48^. The correlation histogram *g(r)* was calculated for binned distances with a bin with of 10 nm and a bandwidth of 5 nm. The data of all STED images per sample were averaged.

In order to compare the pair correlation of real STED images with a simulation of randomly distributed objects, we simulated images comparable to the real data. Using MATLAB (R2019a), 500×500 pixel images with randomly distributed object centers were created. The number of object points per image was selected randomly between 300 and 600 per image and the pixel value was adjusted to 4 (photons/pxl) to match the real data average. Next, the spots were filtered using a 2D Gaussian smoothing kernel with standard deviation of *σ* = 2.5 pixel. Subsequently, 20 simulated images were analyzed using the Huygens Object Analyzer similar to the real data (compare previous section) and finally the pair correlation *g(r)* of simulated data was calculated.

### FCS measurements

For sample preparation see method section Cell culture, transfections and ligand incubation.

#### Calibrations

Calibration of the LSM setup was performed according to established procedures in our research group^49^. Briefly, the optimal correction collar setting was found by minimizing the number of Rhodamine 110 (#83695, Sigma-Aldrich) molecules in the focus. For all our experiments the correction collar matched our coverslip thickness (170 µm). The instrument response function (IRF) was measured using a mirror to enable time-correlated-single-photon-counting (TCSPC) analyses. Next, we measure a Rhodamine 110 solution with 1-5 molecules in the focus to obtain 1) a calibration for the confocal spot shape factor, ***z***_**0**_/***ω***_**0**_ or ***κ***, 2) the ratio of the parallel and perpendicular detection efficiencies, ***γ***, 3) the number and brightness of Rhodamine 110 molecules in the focus and 4) the confocal detection volume by inserting a Rhodamine 110 diffusion constant *D* = 430 µm^²^/s when the calibration was recorded at room temperature (22.5 °C)^50^ or 600 µm²/s when it was recorded at 37 °C considering the temperature dependence of ***D***.

The laser power was measured at the sample using an immersion S170C power meter head (Thorlabs GmbH, Lübeck, Germany) attached to a PM400 power meter body (Thorlabs GmbH, Lübeck, Germany). As the power varied by ∼10% when translating in *x, y* and *z*, we avoid a systematic error by varying the position until maximum power is reached.

#### Recording procedure

A confocal microscope was used to bring the bottom membrane in focus. The diffraction limited focus was placed in a stationary position away from the edge of the cell and away from the endoplasmic reticulum (ER) and Golgi apparatus. FCS curves were recorded during 5 minutes using a 5 µW 488 nm pulsed excitation beam, a 200 µm or 2.1 AU pinhole, a 60X water objective and polarization sensitized readout (see Microscope setups – Confocal setup (‘LSM’)). Solution measurements were performed using identical settings except for placing the focus 50 µm above the glass surface and recording Rhodamine 110 and mEGFP for 1 minute and 5 minutes respectively.

#### FCS curve fitting

All cell measurements were fitted with two diffusion terms, corresponding to a cytoplasmic (cp) and a membrane (mem) component:

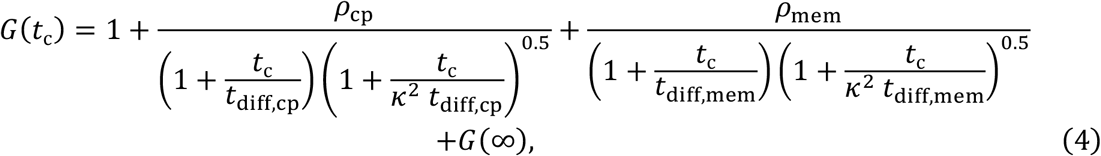

Where *ρ* denotes the species correlation amplitude, *t*_ñrxx_ the species diffusion time, *G(*∞) the residual correlation at infinity, *κ*^6^ the aspect ratio of the focus and *t* the correlation time. As the signal-to-noise was limited, the stability of the fit was improved by not fitting an additional bunching term to account for triplet as it did not affect the values of the diffusion times. To improve the stability of the fit further, a covariance between *t*_ñrxx,îa_ and *t*_ñrxx,o’o_, *t*_ñrxx,îa_was fitted globally over a set of 11 points from 7 CD95 transfected cells, yielding a diffusion time of 0.60 ms to be kept fixed for all subsequent analyses. For more information on obtaining robust results from noisy live cell FCS data see Supplementary Note 2.

Curve weighting according to *σ*_ùû_^51^ was preferred because of its ability to provide accurate weights at long correlation times. Our measurements fulfilled the requirement for that the recording can be divided in >10 chunks of 20 seconds each. FCS curves were created and fitted using the SymPhoTime software (PicoQuant GmbH, Berlin, Germany).

### Confocal Photo Bleaching Step Analysis (cPBSA)

cPBSA measurements were performed on the Abberior setup (compare *Microscope setups*) using circular polarized light and a 100XO objective (NA 1.4, UPLSAPO, Olympus). Since the cell fixation which was needed to immobilize the receptors leads to a deflation of the cell, we ensured that a single membrane layer was in focus by measuring the area underneath the nucleus (see Supplementary Figure 7).

#### Automated data acquisition script

Data acquisition using a confocal microscope is generally slower than TIRF-based PBSA because only one molecular assembly can be measured simultaneously. To gather sufficient statistics, a data acquisition script was written that automates data acquisition after a manual area selection. The program uses the Python Application Programming Interface (API) from the Imspector acquisition software and contains a graphical user interface (GUI). Source code is available on request. Data acquisition works as follows:

1. A suitable area (20 × 20 µm^2^) is selected on the lower membrane by the user.
2. An overview image is recorded using 50nm pixel size, 10 µs dwell time, 5% 488 nm excitation and summed over 3 frames. The output corresponding to 5% laser power fluctuated around 1.3 µW (see Supplementary Table 2).
3. The overview image is smoothened using a Gaussian filter with a standard deviation (sigma) of 1 pixel.
4. Molecular assemblies are identified from local maxima that exceed 3-5 counts on the smoothed image. The threshold level was adjusted per area as needed to select all spots while avoiding crowding by visual inspection.
5. Local maxima that are closer than 450 nm to any other local maxima are not considered for further analysis.
6. A photon trace is recorded for each remaining local maximum by placing the confocal beam there for a duration of 3 seconds.
7. A quick display is rendered for user feedback.

#### Data quality optimization

We established an experimental procedure to optimize the quality of our data. Firstly, our sample fixation procedure minimizes autofluorescence. Secondly, only molecular assemblies that are below the nucleus were recorded to ensure that the lower membrane was not in close proximity to the top membrane, as cells deflate upon fixation (see Supplementary Figure 7). To avoid deflation as far as possible, we forgo upside-down mounting on a cover slip and image cells in well slides instead. Thirdly, low excitation power and integration time was used for creating an overview image in order to avoid premature bleaching.

#### Data analysis

Data analysis was done using the Kalafut-Visscher (KV) algorithm^19^ implemented by Hummert et al. in python^18^. The KV algorithm takes a minimal step size as a sole user input, limiting user bias. As our TCSPC modality records the arrival time of each photon, we can set the time binning of our data (*t*_bin_(s)) post-acquisition. Due to the inherent noise level and varying fluorophore brightness a low threshold will count noise as events, overestimating the real number of Fluorophores, whereas a high threshold will discard bleaching events, underestimating the real number of Fluorophores. The threshold was chosen carefully to balance these two effects at 50 counts per *t*_bin_ of 5 ms, corresponding to 10 kHz at 1.36 µW. To compensate for variations in the laser power, the minimum step size was corrected according to:

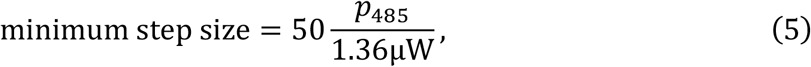

where *p*_485_ is the laser power of the 485 excitation laser for that measurement in µW (see also Supplementary Table 2). Bleaching traces where no steps were found are disregarded from further analysis. No other selection criteria were applied.

#### Fluorescence polarization on traces

Intensities were calculated for traces that showed a single step by integrating fluorescence while the fluorophore was on. As circular polarization was used, fluorescence polarization was calculated using

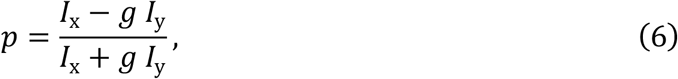

where *I*_x_ is the signal oriented along the x-axis and *I*_y_ was the signal along the y-axis and

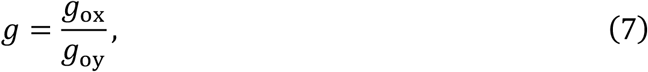

the relative detection efficiency along the x and y axis under circular polarization.

#### Cross-correlation of traces

The *I*_x_ and *I*_y_ signals of detector x- and y-polarization sensitized detectors were cross-correlated and analyzed using the home-built program Kristina^52^. All traces were used without any filtering. The signal-to-noise ratio was very high despite having a low total amount of photons as all photons correlate. Similar to FCS data, the correlation curve was fitted with one diffusion term and 3 bunching terms.

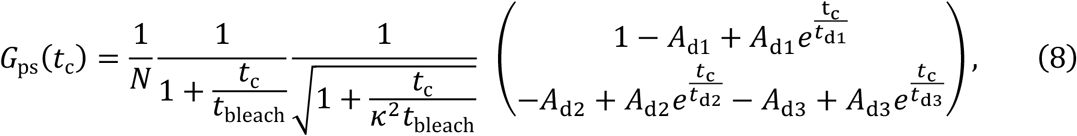

where *κ*^2^was fixed to 100 such that the expression under the root is ∼1, *A* indicate amplitudes, *t* correlation times. Results are summarized in Table 4. While the cross-correlation of a bleaching event is different from a diffusion event, no specialized model for this scenario was available. The resulting residuals around the bleaching time are acceptable as we are mainly interested in the bunching terms. The predicted variance from the cPBSA cross-correlation is discussed in Supplementary Note 3.

### Cell Lifetime FRET Image Spectroscopy (CELFIS)

The method is described in detail in ^53^. First, we obtained the normalized donor only decay,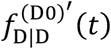, from the donor emission upon donor excitation for a donor only reference sample, 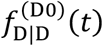:^22^

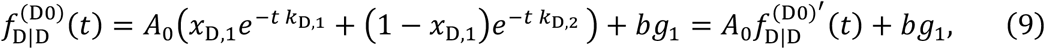

where, *x*_D,i_ and *k*_D,i_ represent the fraction and decay rates of two fluorescence species, *A*_Ñ_ represents the amplitude and *bg*_*i*_ represents the noise floor. This is subsequently used to fit the additional decay of the donor emission upon donor excitation for the donor-acceptor sample^22^

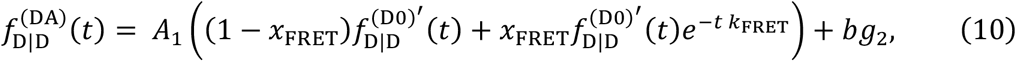

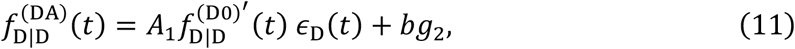

where *x*_FRET_ and *k*_FRET_ are the FRET fraction and FRET rate and we substituted

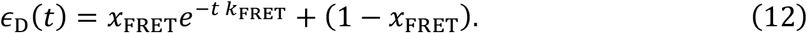

All decays were tail fitted from 1.92 ns to 22.4 ns. Concentrations were determined using a molecular brightness of 814 and 264 Hz / molecule / µW for mEGFP and mCherry, respectively and a maturation factor of 0.8 for both mEGFP and mCherry. The oligomer fraction was obtained from *x*_âäàã_ by calculating the FRET signal corresponding to a pure dimer, *x*_âäàã,opq_. To calculate the latter, we used 1) AV simulations were done in the program Olga^23^ assuming a 51 amino acid linker and effective FRET range up to 80 Å using a solution NMR model of trimeric CD95 TM-domains (pdb id: 2NA7^26^) to set the anchor points for all structures. 2) a 78% and 71% abundance of the heterodimers compared to homodimers for CD95 variants and CD86 on one hand and CTLA4 on the other hand, derived from the abundance of mEGFP and mCherry. 3) an estimated maturation factor of 80%^16,20^.

### Microscope setups

#### Olympus IX83 widefield system

The IX83 P2ZF inverted epi-fluorescence microscope system (Olympus Europa SE & CO. KG, Hamburg, Germany) was used for all widefield and time-laps measurements. The microscope is equipped with the motorized TANGO Desktop stage (Märzhäuser Wetzlar GmbH & Co. KG, Wetzlar, Germany) and the Photometrics Prime BSI camera (Teledyne Photometrics, Tucson, Arizona, USA). An internal halogen lamp and the SOLA Light Engine (Lumencor Inc., Beaverton, Oregon, USA) served as light source for transmitted (brightfield, phase contrast) and reflected (fluorescence) illumination, respectively.

#### Abberior Expert Line setup

STED, cPBSA and FRET measurements were performed on an Abberior Expert Line system as described previously^54^ (Abberior Instruments GmbH, Göttingen, Germany). Additionally, Polarization control for PBSA measurements was achieved using λ/2 and λ/4 waveplates (Abberior Instruments) and a SK010PA-vis 450-800 nm polarization analyzer (Schäfter Kirchhoff GmbH, Hamburg, Germany). Cells were kept at 37 °C using a Heating Insert HP-LabTek (Pecon GmbH, Erbach, Germany). The instrument is operated using the customized Abberior microscope software Imspector (version 14.0.3060, Abberior Instruments GmbH).

#### Confocal setup (‘LSM’)

Fluorescence Correlation Spectroscopy data was recorded using a confocal microscope modified with pulsed excitation and polarization-sensitized time correlated single photon counting readout. Excitation light was created using a Sepia II (PicoQuant GmbH, Berlin, Germany) driving an LDH-D-C-485 laser head (PicoQuant) and coupled to a FluoView1000 IX81 inverted microscope (Olympus, Shinjuku, Japan). Light was focused to a diffraction limited spot using an 60x water immersion UPLSAPO 1.2 NA objective (Olympus) and emitted light was separated using a DM405/488/559/635 quadband mirror (Olympus). Emitted fluorescence was split into perpendicular and parallel components using a polarizing beam splitter and measured using a BrightLine Fluorescence Filter 520/35 (Semrock Inc., Rochester, New York, USA) and PDM series avalanche photo diodes (Micro Photon Devices, Bolzano, Italy) for each channel. Electronic pulses were converted to photon events using a HydraHarp (PicoQuant). Cells were kept at 37 °C using a Heating Insert HP-LabTek (Pecon GmbH, Erbach, Germany).

## Data availability

All data obtained in the study are available from the corresponding authors upon reasonable request.

## Code availability

Code for PBSA trace segment variance prediction is included as a Supplementary Code. All other algorithms have been previously described elsewhere and are correspondingly cited. Analysis notebooks (python files, jupyter notebooks and MATLAB files) are available on reasonable request.

## Author contributions

C.M. and C.A.M.S. conceived and supervised the project. N.B. and N.v.d.V. designed and performed the experiments and analyses. A.B. and C.W. organized and performed Flow Cytometry Measurements, A.G. contributed to the design of plasmids and provided substantial advice for FRET measurements. N.B., N.v.d.V., C.M. and C.A.M.S. discussed the results and wrote the manuscript.

## Acknowledgement

This work was funded by the Deutsche Forschungsgemeinschaft (DFG, German Research Foundation) – project number 267205415 - CRC 1208 (projects A08 and A12). We thank Joél Baudouin (formerly IBS, Grenoble) for providing plasmids and stable cell lines. We thank Sebastian Hänsch and Stefanie Weidtkamp-Peters of the Centre of Advanced Imaging at HHU for fruitful discussions. We thank Suren Felekyan for support with cPBSA and general data analysis and Ralf Kühnemuth for technical support. We thank Hadeel Khalouf and Xiaoyue Shang for cloning initiatives of the pseudo-dimer. We are also in debt to Wolfgang Schulz, and Michèle Hoffmann both in the department of Urology, Heinrich Heine University, for the use of the MACSQuant Analyzer 10. A.G. is thankful for financial support from the HHU Stategischer Forschungsfonds. A.B. is thankful for financial support from the Medical Research School, DSO, Heinrich-Heine-University, Düsseldorf. C.M. is grateful for financial support by VolkswagenFoundation and Fonds der Chemischen Industrie

## References

1 Aggarwal, B. B. Signalling pathways of the TNF superfamily: A double-edged sword. Nature Reviews Immunology 3, 745–756 (2003). https://doi.org:10.1038/nri1184

2 Locksley, R. M., Killeen, N. & Lenardo, M. J. The TNF and TNF receptor superfamilies: Integrating mammalian biology. Cell 104, 487–501 (2001). https://doi.org:10.1016/s0092-8674(01)00237-9

3 Bodmer, J. L., Schneider, P. & Tschopp, J. The molecular architecture of the TNF superfamily. Trends in Biochemical Sciences 27, 19–26 (2002). https://doi.org:10.1016/s0968-0004(01)01995-8

4 Bremer, E. Targeting of the tumor necrosis factor receptor superfamily for cancer immunotherapy. ISRN Oncol 2013, 371854 (2013). https://doi.org:10.1155/2013/371854

5 Chan, F. K. M. Three is better than one: Pre-ligand receptor assembly in the regulation of TNF receptor signaling. Cytokine 37, 101–107 (2007). https://doi.org:10.1016/j.cyto.2007.03.005

6 Chan, F. K. M. et al. A domain in TNF receptors that mediates ligand-independent receptor assembly and signaling. Science 288, 2351–2354 (2000). https://doi.org:10.1126/science.288.5475.2351

7 Boschert, V. et al. Single chain TNF derivatives with individually mutated receptor binding sites reveal differential stoichiometry of ligand receptor complex formation for TNFR1 and TNFR2. Cellular Signalling 22, 1088–1096 (2010). https://doi.org:10.1016/j.cellsig.2010.02.011

8 Naismith, J. H., Devine, T. Q., Brandhuber, B. J. & Sprang, S. R. Crystallographic Evidence for Dimerization of Unliganded Tumor Necrosis Factor Receptor. Journal of Biological Chemistry 270, 13303–13307 (1995). https://doi.org:10.1074/jbc.270.22.13303

9 Vanamee, E. S. & Faustman, D. L. Structural principles of tumor necrosis factor superfamily signaling. Science Signaling 11, eaao4910 (2018). https://doi.org:10.1126/scisignal.aao4910

10 Vanamee, E. S., Lippner, G. & Faustman, D. L. Signal Amplification in Highly Ordered Networks Is Driven by Geometry. Cells 11, 272 (2022). https://doi.org:10.3390/cells11020272

11 Siegel, R. M. et al. SPOTS: signaling protein oligomeric transduction structures are early mediators of death receptor-induced apoptosis at the plasma membrane. Journal of Cell Biology 167, 735–744 (2004). https://doi.org:10.1083/jcb.200406101

12 Weidtkamp-Peters, S. et al. Multiparameter fluorescence image spectroscopy to study molecular interactions. Photochemical & Photobiological Sciences 8, 470–480 (2009). https://doi.org:10.1039/b903245m

13 Lerner, E. et al. FRET-based dynamic structural biology: Challenges, perspectives and an appeal for open-science practices. Elife 10, e60416 (2021).

14 Gerken, M. et al. Fluorescence correlation spectroscopy reveals topological segregation of the two tumor necrosis factor membrane receptors. Biochimica Et Biophysica Acta-Biomembranes 1798, 1081–1089 (2010). https://doi.org:10.1016/j.bbamem.2010.02.021

15 Jaqaman, K., Galbraith, J. A., Davidson, M. W. & Galbraith, C. G. Changes in single-molecule integrin dynamics linked to local cellular behavior. Molecular Biology of the Cell 27, 1561–1569 (2016). https://doi.org:10.1091/mbc.E16-01-0018

16 Ulbrich, M. H. & Isacoff, E. Y. Subunit counting in membrane-bound proteins. Nature Methods 4, 319–321 (2007). https://doi.org:10.1038/Nmeth1024

17 Clarke, S. et al. Covalent Monofunctionalization of Peptide-Coated Quantum Dots for Single-Molecule Assays. Nano Letters 10, 2147–2154 (2010). https://doi.org:10.1021/nl100825n

18 Hummert, J. et al. Photobleaching step analysis for robust determination of protein complex stoichiometries. Molecular Biology of the Cell 32, ar35 (2021). https://doi.org:10.1091/mbc.E20-09-0568

19 Kalafut, B. & Visscher, K. An objective, model-independent method for detection of non-uniform steps in noisy signals. Computer Physics Communications 179, 716–723 (2008). https://doi.org:10.1016/j.cpc.2008.06.008

20 Dunsing, V. et al. Optimal fluorescent protein tags for quantifying protein oligomerization in living cells. Scientific reports 8, 1–12 (2018).

21 Greife, A. et al. Structural assemblies of the di- and oligomeric G-protein coupled receptor TGR5 in live cells: an MFIS-FRET and integrative modelling study. Scientific Reports 6, 36792 (2016). https://doi.org:10.1038/srep36792

22 Peulen, T.-O., Opanasyuk, O. & Seidel, C. A. Combining graphical and analytical methods with molecular simulations to analyze time-resolved FRET measurements of labeled macromolecules accurately. The Journal of Physical Chemistry B 121, 8211–8241 (2017).

23 Dimura, M. et al. Automated and optimally FRET-assisted structural modeling. Nature Communications 11, 5394 (2020). https://doi.org:10.1038/s41467-020-19023-1

24 Clayton, A. H. A. & Chattopadhyay, A. Taking Care of Bystander FRET in a Crowded Cell Membrane Environment. Biophysical Journal 106, 1227–1228 (2014). https://doi.org:10.1016/j.bpj.2014.02.004

25 Dimura, M. et al. Quantitative FRET studies and integrative modeling unravel the structure and dynamics of biomolecular systems. Current opinion in structural biology 40, 163–185 (2016).

26 Fu, Q. S. et al. Structural Basis and Functional Role of Intramembrane Trimerization of the Fas/CD95 Death Receptor. Molecular Cell 61, 602–613 (2016). https://doi.org:10.1016/j.molcel.2016.01.009

27 Karathanasis, C. et al. Single-molecule imaging reveals the oligomeric state of functional TNF alpha-induced plasma membrane TNFR1 clusters in cells. Science Signaling 13, eaax5647 (2020). https://doi.org:10.1126/scisignal.aax5647

28 Rodseth, L. E. et al. 2 CRYSTAL FORMS OF THE EXTRACELLULAR DOMAIN OF TYPE-I TUMOR-NECROSIS-FACTOR RECEPTOR. Journal of Molecular Biology 239, 332–335 (1994). https://doi.org:10.1006/jmbi.1994.1371

29 Papoff, G. et al. Identification and characterization of a ligand-independent oligomerization domain in the extracellular region of the CD95 death receptor. Journal of Biological Chemistry 274, 38241–38250 (1999). https://doi.org:10.1074/jbc.274.53.38241

30 Siegel, R. M. et al. Fas preassociation required for apoptosis signaling and dominant inhibition by pathogenic mutations. Science 288, 2354–2357 (2000). https://doi.org:10.1126/science.288.5475.2354

31 Frey, L., Lakomek, N. A., Riek, R. & Bibow, S. Micelles, Bicelles, and Nanodiscs: Comparing the Impact of Membrane Mimetics on Membrane Protein Backbone Dynamics. Angew Chem Int Edit 56, 380–383 (2017). https://doi.org:10.1002/anie.201608246

32 Liesche, C. et al. CD95 receptor activation by ligand-induced trimerization is independent of its partial pre-ligand assembly. bioRxiv, 293530 (2018). https://doi.org:10.1101/293530

33 Banner, D. W. et al. Crystal structure of the soluble human 55 kd TNF receptor-human TNFβ complex: Implications for TNF receptor activation. Cell 73, 431–445 (1993). https://doi.org:10.1016/0092-8674(93)90132-a

34 Gülcüler Balta, G. S. et al. 3D Cellular Architecture Modulates Tyrosine Kinase Activity, Thereby Switching CD95-Mediated Apoptosis to Survival. Cell Reports 29, 2295–2306 (2019). https://doi.org:10.1016/j.celrep.2019.10.054

35 Henkler, F. et al. The extracellular domains of FasL and Fas are sufficient for the formation of supramolecular FasL-Fas clusters of high stability. Journal of Cell Biology 168, 1087–1098 (2005). https://doi.org:10.1083/jcb.200501048

36 Scott, F. L. et al. The Fas-FADD death domain complex structure unravels signalling by receptor clustering. Nature 457, 1019–1022 (2009). https://doi.org:10.1038/nature07606

37 Esposito, D. et al. Solution NMR Investigation of the CD95/FADD Homotypic Death Domain Complex Suggests Lack of Engagement of the CD95 C Terminus. Structure 18, 1378–1390 (2010). https://doi.org:10.1016/j.str.2010.08.006

38 Levoin, N., Jean, M. & Legembre, P. CD95 Structure, Aggregation and Cell Signaling. Frontiers in Cell and Developmental Biology 8, 314 (2020). https://doi.org:10.3389/fcell.2020.00314

39 Wang, L. W. et al. The Fas-FADD death domain complex structure reveals the basis of DISC assembly and disease mutations. Nature Structural & Molecular Biology 17, 1324–U1176 (2010). https://doi.org:10.1038/nsmb.1920

40 Potma, E. O. et al. Reduced protein diffusion rate by cytoskeleton in vegetative and polarized Dictyostelium cells. Biophysical Journal 81, 2010–2019 (2001). https://doi.org:10.1016/s0006-3495(01)75851-1

41 Reis, C. R., van Assen, A. H. G., Quax, W. J. & Cool, R. H. Unraveling the Binding Mechanism of Trivalent Tumor Necrosis Factor Ligands and Their Receptors. Molecular & Cellular Proteomics 10, M110.002808 (2011). https://doi.org:10.1074/mcp.M110.002808

42 Berger, R. M. L. et al. Nanoscale FasL Organization on DNA Origami to Decipher Apoptosis Signal Activation in Cells. Small 17, 2101678 (2021). https://doi.org:10.1002/smll.202101678

43 Liesche, C. et al. Death receptor-based enrichment of Cas9-expressing cells. Bmc Biotechnology 16, 17 (2016). https://doi.org:10.1186/s12896-016-0250-4

44 Fricke, F., Beaudouin, J., Eils, R. & Heilemann, M. One, two or three? Probing the stoichiometry of membrane proteins by single-molecule localization microscopy. Scientific Reports 5, 14072 (2015). https://doi.org:10.1038/srep14072

45 Qureshi, O. S. et al. Constitutive Clathrin-mediated Endocytosis of CTLA-4 Persists during T Cell Activation. Journal of Biological Chemistry 287, 9429–9440 (2012). https://doi.org:10.1074/jbc.M111.304329

46 Schindelin, J. et al. Fiji: an open-source platform for biological-image analysis. Nature Methods 9, 676–682 (2012). https://doi.org:10.1038/nmeth.2019

47 Stoyan, D. & Stoyan, H. Estimating pair correlation functions of planar cluster processes. Biometrical Journal 38, 259–271 (1996). https://doi.org:10.1002/bimj.4710380302

48 Peckys, D. B., Korf, U. & de Jonge, N. Local variations of HER2 dimerization in breast cancer cells discovered by correlative fluorescence and liquid electron microscopy. Science Advances 1, e1500165 (2015). https://doi.org:10.1126/sciadv.1500165

49 Felekyan, S. et al. Full correlation from picoseconds to seconds by time-resolved and time-correlated single photon detection. Review of Scientific Instruments 76, 083104 (2005). https://doi.org:10.1063/1.1946088

50 Gendron, P. O., Avaltroni, F. & Wilkinson, K. J. Diffusion Coefficients of Several Rhodamine Derivatives as Determined by Pulsed Field Gradient-Nuclear Magnetic Resonance and Fluorescence Correlation Spectroscopy. Journal of Fluorescence 18, 1093–1101 (2008). https://doi.org:10.1007/s10895-008-0357-7

51 Wohland, T., Rigler, R. & Vogel, H. The standard deviation in fluorescence correlation spectroscopy. Biophysical Journal 80, 2987–2999 (2001). https://doi.org:10.1016/s0006-3495(01)76264-9

52 Borst, J. W. et al. Structural Changes of Yellow Cameleon Domains Observed by Quantitative FRET Analysis and Polarized Fluorescence Correlation Spectroscopy. Biophysical Journal 95, 5399–5411 (2008). https://doi.org:10.1529/biophysj.107.114587

53 van der Voort, N. T. M., Bartels, N., Monzel, C. & Seidel, C. A. M. Quantifying the Spatio-temporal Evolution of Protein Interactions using Cell Lifetime FRET Image Spectroscopy (CELFIS). bioRxiv (2022).

54 Budde, J.-H. et al. FRET nanoscopy enables seamless imaging of molecular assemblies with sub-nanometer resolution. arXiv, 2108.00024 (2021).

