## Supplementary Information for "A Minimal Model of CD95 Signal Initiation Revealed by Advanced Super-resolution and Multiparametric Fluorescence Microscopy"

† contributed equally

Bibliography ..................................................................... **Fehler! Textmarke nicht definiert.**

### Supplementary Notes

#### Supplementary Note 1: Optimal instrument settings for live cell FCS

For method details see method section *FCS measurements*.

Live cell FCS measurements on mEGFP remain challenging due to the limited mEGFP photostability and limited mEGFP abundance in a cell. To nevertheless obtain a robust readout, we discuss optimal experimental settings along with a brief description of the photophysical effects governing the observations.

**Pinhole setting**

The optimal pinhole setting was determined experimentally to be 200 µm in diameter, or 492 nm backprojected pinhole radius[1], which corresponds to 2.1 Airy Units (AU). This setting optimally balances 1) a high photon collection efficiency 2) a sharp Point Spread Function (PSF) 3) the PSF shape to resemble a Gaussian. The tradeoff consists thereof that an open pinhole with high collection efficiency is needed to compensate for the poor photo-stability of mEGFP and resulting low signal-to-noise ratio (SNR). However, opening the pinhole transforms the shape of the PSF from a $sinc^2$, which is Gaussian-like to a sinc function, which in not Gaussian-like. As FCS theory (see Equation (4)) models a molecule diffusing through a 3D Gaussian volume, an open pinhole results in a mismatch between model and measurement visible in the fit residuals. As reported by others[2], a 2.1 AU pinhole leads to small but acceptable deviations between the model function and data.

**Fluorescent molecule concentration changes during measurement**

A change in fluorescent protein concentration is registered by the correlation function at long time scales, which complicates fitting slow membrane diffusion. For solution measurements, the dominant process for concentration decrease is adsorption of to the glass surface, which is easily prevented by coating the glass surfaces with BSA (incubate 1 mg/ml Bovine Serum Albumin for 10 minutes, BSA, Sigma-Aldrich Merck group, Taufkirchen, Germany). Bleaching does not significantly affect concentrations in solution measurements as the bleaching rate is small compared to the large fluorophore reservoir. In cells, a change in fluorescent protein concentration cannot be circumvented as photo-bleaching can readily deplete the reservoir of fluorescent proteins at an organelle or cellular scale. To mitigate the effects of a decreasing mEGFP concentration on the FCS curve, we divide the photon trace in chunks of approximately constant concentration and average the pieces[2].

We are able to gain additional insight in the photo-bleaching process from synergistically combining our read-out from FCS and cPBSA. From our FCS measurements we obtain diffusion times and fluorophore brightness, which we use to calculate the average number of photons per time the molecule diffuses through the focus to be ~1.5 for mEGFP[3]. From cPBSA we are able to obtain the total photon budget of mEGFP to be ~1000 photons. Taken together we conclude that the probability of mEGFP bleaching during a single pass through the detection volume is very low and that the mEGFP concentration decreases because a single molecule passes through the detection volume many times.

**Power setting & photon budget**

A higher laser power increases the signal-to-noise ratio for the FCS curve at the cost of a higher bleaching rate, which cause unwanted changes in local concentrations (see section above). In this section, we explain the underlying processes and obtain a trade-off between the SNR level of the FCS curve and the bleaching rate.

Primarily, the SNR of an FCS curve must be sufficient to enable interpretation, which scales with the number of photons detected while a single molecule diffuses through the focus[4]. Interestingly, our results indicate that the average number of photons is ~1.5. On the condition that molecules diffuse independent from each other, at least two photons are needed to obtain a correlation. This apparent contradiction is resolved by realizing that the number of photons per event follows a distribution with a long tail at higher photon numbers. I.e., while some of the molecules emit zero or one photon, the fraction which emits two or more photons is responsible for the correlation in FCS[5].

To help understand bleaching processes, we introduce the concept of photon budget to mean the total amount of photons emitted by the fluorophore before bleaching. It is inversely proportional to the bleaching probability per excitation cycle. Work done by others[6-8] on decay pathway modelling reveals that the photon budget of mEGFP is constant at low irradiance but decreases after a transition regime. The decrease in photon budget is due to an additional photon being absorbed while the molecule is in the excited state, opening up additional photo-bleaching pathways and increasing the photo-bleaching probability per cycle. While a laser power lower than the transition irradiance maximizes the photon budget, a definite number was not found in literature by the authors, although an upper limit was reported by Cranfill et al.[7] to be 80 µW using 488 nm excitation in a diffraction limited focus. Based on our own experimental experience we estimate the transition point from mEGFP to be lower than ~10 µW.

To satisfy all the criteria above, the laser power was experimentally determined to be 5 µW corresponding to 3 $kW/cm^2$ for a calibrated 0.165 $\mu m^2$ focal area.

**Recording time**

Longer recording times improve the SNR of the FCS curve. However, to sample sufficient cell-to-cell variation during a measurement day it was limited to 5 minutes.

#### Supplementary Note 2: Live-cell Membrane FCS

To verify that CD95 is sufficiently mobile and hence able to form (higher) oligomers, we determined CD95 diffusion constants D during the whole signaling process using Fluorescence Correlation Spectroscopy (FCS). FCS was performed on live cells for $CD95_{D0}$ (Supplementary Figure 3a) and $CD95(\Delta DD)_{D0}$ (Supplementary Figure 3b) before and 100 - 200 minutes after ligand addition as well as for $CD86_{D0}$ and $CTLA4_{DA}$ as single and double transmembrane helix references, respectively (Supplementary Figure 3c). FCS curves were generated for each cell and fitted with two diffusion terms and no bunching term (see methods). The fast diffusion term was attributed to the presence of cytoplasmic mEGFP, which was confirmed by 3D confocal images of live cells. As the confocal detection volume extends halfway into the cytoplasm, FCS is sensitive to mEGFP present in the cytoplasm (see Supplementary Figure 12). To confirm that the fast diffusion component was of cytoplasmic origin, we fitted the fast diffusion term globally for all curves from the CD95 sample yielding a value of $t_{\mathrm{diff,cp}} = 0.60$ ms (corresponding to $D = 20$ µm$^2$/s), reminiscent of soluble protein diffusion in eukaryotic cells, with typical values of 24 µm$^2$/s[9]. In addition, we measured diffusion of free mEGFP in the cytoplasm and obtained two diffusion times, 0.27 ms and 2.2 ms, with a weighted average time of 0.50 ms (see Supplementary Figure 4) close to the cytoplasmic component of CD95. In the following, the fast diffusion term was kept fixed for all samples to improve the sensitivity of the fit for the slow diffusion time (see Supplementary Figure 3a-c). The time of the slow diffusion process turned out to lie in the range of 30-100ms and matches literature values for membrane proteins[10,11]. No difference in the diffusion constant was found between CD95 species within the measurement accuracy (see Supplementary Figure 3d), with absolute values of $D = 0.21$-$0.24$ µm$^2$/s. Interestingly, $CTLA4_{DA}$ showed similar diffusion times of $D = 0.19$ µm$^2$/s, but CD86 showed significantly slower diffusion times of $D = 0.15$ µm$^2$/s indicating that the number of transmembrane helices is not the dominant factor of receptor diffusion. This result indicates that CD95 is sufficiently mobile to exhibit dynamic changes in its oligomeric state over time. In addition, comparison of the absolute values suggests that CD95 does not form supramolecular structures as this would result in a highly decreased diffusion constant. Finally, we tested whether the mobility of CD95 would change after ligand addition. To this end, CD95 and CD95(ΔDD) diffusion was monitored over 100-200 minutes after ligand addition (see Supplementary Figure 3e). Overall, our data confirm sustained CD95 mobility during the whole signaling process. Despite this possibility to accumulate into higher ordered structures CD95 did not show any systematic change in CD95 diffusion, thus indicating no excessive change in the receptor oligomerization state.

#### Supplementary Note 3: Predicted variance from cPBSA cross-correlation curves.

The cross-correlation of cPBSA traces is described in the method section on cPBSA.

To predict the variance, we assume a simple model where the mEGFP molecule can enter a dark state with rate $k_{\mathrm{off}}$ and return to the bright state with rate $k_{\mathrm{on}}$. The latter can directly be obtained from the cross correlation fit as it is inversely proportional to $t_{\mathrm{on}}$, the characteristic time from the cross-correlation bunching therm. To obtain $k_{\mathrm{off}}$, we define the fraction of time spent in the on state, $\alpha$ as

$$\alpha = \frac{k_{\mathrm{on}}}{k_{\mathrm{on}} + k_{\mathrm{off}}}, \tag{S1}$$

Where $(1 - \alpha)$ equals the amplitude of the bunching therm. Using the values from the fit listed in Supplementary Table 4, we obtain 1/5 ms and 1/30 ms for $k_{\mathrm{on}}$ and $k_{\mathrm{off}}$ for the slowest transition. To obtain a theoretical description of the variance on a trace segment, we define

$$n = t_{\mathrm{bin}}(k_{\mathrm{on}} + k_{\mathrm{off}}), \tag{S2}$$

where $n$ is the average number of transitions in some time period $t_{\mathrm{bin}}$. For a fluorophore with fluorescence rate $k_{\mathrm{fl}}$, the average signal in time period $t_{bin}$ is given by

$$\langle S \rangle = t_{\mathrm{bin}} \langle k_{\mathrm{fl}} \rangle \alpha; \tag{S3}$$

$$\langle S \rangle = N_{\mathrm{fl}} \alpha, \tag{S4}$$

where $N_{\mathrm{fl}}$ is defined as the average number of fluorophores emitted if there was no dark state. Note that the fluorophore brightness, $k_{\mathrm{fl}}$ is understood to include polarization effects, such that it differs per molecule. We may now write the variance of the signal as

$$Var(S) = \left|\frac{\partial S}{\partial \alpha}\right|^2 Var(\alpha) + \left|\frac{\partial S}{\partial N_{\mathrm{fl}}}\right|^2 Var(N_{\mathrm{fl}}), \tag{S5}$$

where the covariance between $\alpha$ and $N_{\mathrm{fl}}$ as a function of excitation power is not considered as the excitation was kept within the range 1.3 ± 0.2μW (Supplementary Table 2). From Barth et al.[12], we obtain the variance of alpha

$$Var(\alpha) = \alpha(1 - \alpha)\frac{2}{n}\left(1 + \frac{e^{-n} - 1}{n}\right) \tag{S6}$$

To obtain the variance of $N_{\mathrm{fl}}$, we consider that the average number of photons emitted follows a Poisson distribution

$$Var(N_{\mathrm{fl}}) = \frac{N_{\mathrm{fl}}}{\alpha}. \tag{S7}$$

Combining all formulas, we obtain a direct expression:

$$Var(S) = N_{\mathrm{fl}}^2 \alpha(1 - \alpha)\frac{2}{n}\left(1 + \frac{e^{-n} - 1}{n}\right) + \alpha N_{\mathrm{fl}}. \tag{S8}$$

We check that in the limit of very fast fluctuations from the dark state, the variance due to $\alpha$ becomes zero and we obtain a Poisson distribution as expected

$$\lim_{n\to\infty} Var(S) = \alpha N_{\mathrm{fl}} \qquad (S9)$$

Intuitively, we understand that long dark state times with respect to $t_{\mathrm{bin}}$ cause fluctuations in $\alpha$ whose variance dominates the inherent Poisson noise, hence this result confirms that our initial model of considering only the longest dark states captures all essential features. To check our expression, we perform Monte Carlo simulations for all values of $\alpha$, $n$ and $N_{\mathrm{fl}}$ (see Supplementary Figure 11 and Supplementary Code 1) confirming that it correctly predicts the variance and showing that the variance per trace fluctuates with the stochastic number of blinks.

**Supplementary Note 4:** To investigate the possibility of higher oligomeric states further, we focus our attention on bright areas on the membrane of typically ~1µm diameter and ~5µm separation visible for all constructs including controls which cannot be related to intracellular organelles in close membrane proximity or membrane ruffling (Figure 5b and Supplementary Figure 1). We probe the oligomeric state of the bright areas by modifying our analysis to only consider signal originating from there, yielding a comparable although slightly higher oligomerization state from before, which is expected due to concentration driven kinetics (see (N.v.d.V., N.B., C.M., C.A.M.S., *manuscript in preparation*)). As the bright areas are not specific to CD95 variants and the FRET signature is similar before, we conclude that the bright areas are not higher oligomeric states, but simply local concentration of receptors ubiquitous to membrane receptors.

### Supplementary Tables

| Sample | Statistics [cells] | Membrane fraction | Cytoplasmic fraction |
|---|---|---|---|
| **CD86$_{DO}$** | 9 | 0.589 ± 0.085 | 0.411 ± 0.085 |
| **CD86$_{DO}$** | 6 | 0.628 ± 0.052 | 0.372 ± 0.052 |
| **CTLA4$_{DA}$** | 10 | 0.703 ± 0.036 | 0.297 ± 0.036 |
| **CD95$_{D0}$** | 11 | 0.594 ± 0.045 | 0.406 ± 0.045 |
| **CD95(ΔDD)$_{D0}$** | 12 | 0.598 ± 0.052 | 0.402 ± 0.052 |
| **CD95$_{D0}$ + Lig** | 11 | 0.597 ± 0.043 | 0.403 ± 0.043 |
| **CD95(ΔDD)$_{D0}$ + Lig** | 14 | 0.637 ± 0.030 | 0.363 ± 0.030 |
| **Total** | 73 | 0.621 ± 0.062 | 0.379 ± 0.062 |

**Supplementary Table 1: Membrane and cytoplasmic fraction of cell fluorescence signal determined with FCS**

Fractions of fast cytoplasmic and slow membrane diffusion for different membrane proteins measured with FCS. See Supplementary Note 1 and Supplementary Figure 3.

| dataset | date | laser power [µW] | minimum step size [counts] |
|---|---|---|---|
| **CD95** | 22 July 2021 | 1.36 | 50 |
| **CD95 +L** | 23 July 2021 | 1.60 | 58 |
| **CD95(ΔDD)** | 22 July 2021 | 1.36 | 50 |
| **CD95(ΔDD) +L** | 23 July 2021 | 1.60 | 58 |
| **CD86-mEGFP$_1$** | 22 July 2021 | 1.36 | 50 |
| **CD86-mEGFP$_2$** | 24 November 2021 | 1.37 | 50 |
| **CTLA4$_{DA}$** | 2 February 2022 | 1.00 | 36 |
| **CD86-mEGFP-mEGFP** | 2 February 2022 | 1.00 | 36 |

**Supplementary Table 2: Laser power fluctuations during measurement days for PBSA data.**

The step threshold was adjusted such that the ratio of the power and the threshold remains constant (see Supplementary Figure 8).

| model No. therms | $\chi^2_{red,avg}$ | $t_{diff,1}$ | $t_{bunch,1}$ | $t_{bunch,2}$ | $t_{bunch,3}$ | $t_{bunch,4}$ |
|---|---|---|---|---|---|---|
| **1 diffusion, 1 bunching[1]** | 6.7 | 59.6 ms | 0.85 ms | - | - | - |
| **1 diffusion, 2 bunching[1]** | 2.1 | 73 ms | 2.41 ms | 0.09 ms | - | - |
| **1 diffusion, 3 bunching** | 1.08 | 83 ms | 5 ms | 0.56 ms | 0.011 ms | - |
| **1 diffusion, 4 bunching[2]** | 0.78 | 95 ms | 13.4 ms | 1.4 ms | 0.11 ms | 0.006 ms |

**Supplementary Table 3: mEGFP bunching terms.**

The correct model for fitting cross correlation of cPBSA traces was determined by best $\chi^2_{\text{red,avg}}$, the reduced chi-square averaged over the four cross correlations shown in Supplementary Figure 10a. Correlation times were fitted globally over the four cross correlations, the amplitudes were fitted individually. [1]Fit has too high $\chi^2_{\text{red}}$ to describe data well. [2] $\chi^2_{\text{red}} < 1$ indicates overfitting.

| Parameter | CD95 | CD95 + L | CD95(ΔDD) | CD86 |
|---|---|---|---|---|
| **$G(\infty)$** | 0.95 | 1.01 | 0.95 | 0.98 |
| **N** | 0.78 | 0.72 | 0.98 | 0.77 |
| **$t_{bleach}$ [ms]*** | 84 | 84 | 84 | 84 |
| **$A_{bleach}$ [%]**** | 69.9 | 68.5 | 68.5 | 64.9 |
| **κ *** | 100 | 100 | 100 | 100 |
| **$A_{d1}$ [%]** | 13.3 | 14.0 | 11.0 | 15.6 |
| **$t_{d1}$ [ms]*** | 5.0 | 5.0 | 5.0 | 5.0 |
| **$A_{d2}$ [%]** | 10.0 | 11.5 | 13.0 | 11.7 |
| **$t_{d2}$ [ms]*** | 0.56 | 0.56 | 0.56 | 0.56 |
| **$A_{d3}$ [%]** | 6.8 | 6.0 | 7.5 | 7.8 |
| **$t_{d3}$ [ms]*** | 0.011 | 0.011 | 0.011 | 0.011 |

**Supplementary Table 4: Fit parameters for cross-correlation fits on ensemble traces** (see Supplementary Figure 10 and Equation (8)). *fitted globally ** calculated using $A_{\mathrm{bleach}} = 100 - A_{\mathrm{d1}} - A_{\mathrm{d2}} - A_{\mathrm{d3}}$.

| plasmid id | sequence | linker sequence | flexible part protein #aa | linker length #aa | flexible part FP #aa | Total flexible #aa |
|---|---|---|---|---|---|---|
| **1714** | CD95(1-335)-mCherry T2A CD95(1-335)-mEGFP | GGGGGPVPQWEGFAALLATPVAT/ GGGGGPVPQWEGFAALLATPVGGAV | 9[1] | 23 / 25 | 16 / 12 | 48 / 46 |
| **1695** | CD95-ΔDD(1-210)-mCherry T2A CD95-ΔDD(1-210)-mEGFP | GGGPVPQWEGFAALLATPVAT / GGGPVPQWEGFAALLATPVGGAV | 16[4] | 21 / 23 | 16 / 12 | 53 / 51 |
| **1693** | CTLA4(1-200)-mCherry T2A CTLA4(1-200)-mEGFP | GGGPVPQWEGFAALLATPVAT / GGGPVPQWEGFAALLATPVGGAV | 16[4] | 21 / 23 | 16 / 12 | 53 / 51 |
| **1706** | CD86-mCherry T2A CD86-mEGFP | GGGPVPQWEGFAALLATPVAT / GGGPVPQWEGFAALLATPVGGAV | -[2] | 21 / 23 | 16 / 12 | 37 / 35 |
| **1531** | CD95(1-335)-mEGFP | GGGGGPVPQWEGFAALLATPVGGAV | 9[1] | 25 | 12 | 46 |
| **1516** | CD95-ΔDD(1-210)-mEGFP | GGGPVPQWEGFAALLATPVGGAV | 16 | 23 | 12 | 51 |
| **1706** | CD86-mEGFP | GGGPVPQWEGFAALLATPVGGAV | -[2] | 23 | 12 | 35 |
| **88** | CD86-*link*-mEGFP-mEGFP[3] | GGGPVPQWEGFAALLATPVGGAV | -[2] | 23 | 12 | 35 |
| **88** | CD86-mEGFP-*link*-mEGFP[3] | GSSGSSNAAIINAAGSSGSS | 11 | 20 | 12 | 43 |

**Supplementary Table 5: Specifications of linker lengths for used constructs.**

[1]9 amino acids are used to model the flexible death domain, see methods. [2]flexible part could not be estimated because the structure of the CD86 TM domain is not known, taken as 0. [3]link indicates the position of the linker detailed. #aa: number of amino acids. CTLA-mEGFP control was not used as it did not localize near the membrane. [4] linker length based on the residual 16 intracellular amino acids after transmembrane domain.

### Supplementary Codes

```
#all units in s, 1/s
#this code is fast in c, but slow in python
def simulateTrace(Nfl, alpha, n, tbin = 5e-3, tstep = 1e-4, timestop = 1):
  """
    Do a Monte Carlo simulation of a trace, assuming a single dark state.
    tstep is the time resolution of the simulation. It should be chosen such
  that the probability of multiple blinking events in 1 step is low, i.e.,
  pon and poff < 0.1
  Nfl:    average number of fluorophores in time tbin if the molecule is on,
          i.e. the molecule brightness (dimensionless)
  alpha:   the average fraction of time the molecule spends in the on state (dimensionless)
  n:      the average number of blinks in time tbin (dimensionless)
  tbin:    time period (s)
  tstep:   time resolution of simulation, see above (s)
  timestop: amount of time to simulate (s)"""
  #calculate derived variables
  Nevents = int(np.ceil(timestop / tstep))
  pfl = Nfl / tbin * tstep
  pon = n * alpha / tbin * tstep
  poff = n * (1-alpha) / tbin * tstep
  #fluorophore starts in the on state, compliant with physical conditions.
  state = 'on'
  #initialize arrays
  events = np.zeros(Nevents)
  #event loop
  for i in range(Nevents):
    if state == 'on':
      #add a poissonian number of photons
      events[i] = np.random.poisson(pfl)
      #switch off with probability poff
      if poff > np.random.random():
        state = 'off'
    elif state == 'off':
      #switch on with probability pon
      if pon > np.random.random():
        state = 'on'
  #downsample trace to tbin
  binfact = int(np.ceil(tbin / tstep))
  nbins = int(np.ceil(timestop / tbin))
  trace = np.sum(events.reshape((nbins, binfact)), axis = 1)
  #calculate variance
  variance = np.var(trace)
  return variance, trace
```

**Supplementary Code 1: Code for Monte Carlo simulations on variance predictions.** dark state Monte Carlo simulations for PBSA trace segment variance prediction. Code was tested in python 3.7, but should also run in Python 2.x and 3.x versions. The only dependency is the numpy library. Results are summarized in Figure 11.

### Supplementary Figures

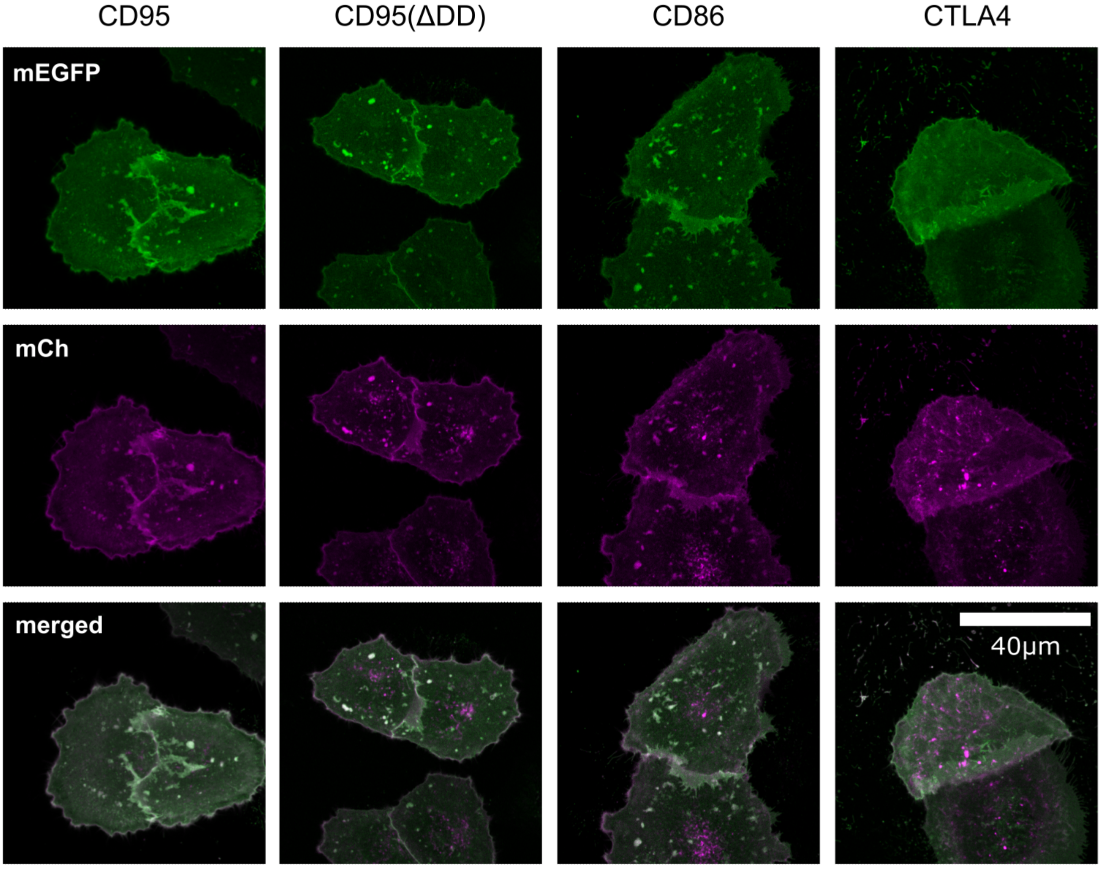

**Supplementary Figure 1: Confocal images of transfected cells show the protein localization in the membrane.**

Images show live Hela CD95$^{KO}$ cells transfected with bicistronic plasmids coding for donor (mEGFP) and acceptor (mCherry) bound CD95, CD95(ΔDD), CD86 or CTLA4 during FRET measurements, focused on the lower cell membrane. Higher intensities at cell edges and cell-to-cell contact points show the correct integration of the membrane proteins into the outer cell membrane. Scale bar applies for all images.

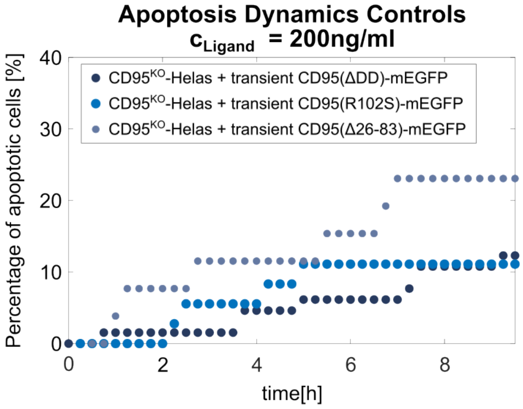

**Supplementary Figure 2: Apoptosis dynamics of CD95 variants.**
Apoptosis Dynamics of transiently transfected Hela $CD95^{KO}$ cells with the CD95 variants CD95(ΔDD), CD95(R102S) and CD95(Δ26-86). While the first two variants show apoptosis caused by natural apoptosis or transfection stress, the PLAD- depleted variant CD95(Δ26-86) (also called CD95(ΔPLAD)) shows an increased apoptosis efficiency up to 25% of dead cells. Statistics: > 25 cells for CD95(R102S) and CD95(Δ26-86), > 65 cells for CD95(ΔDD).

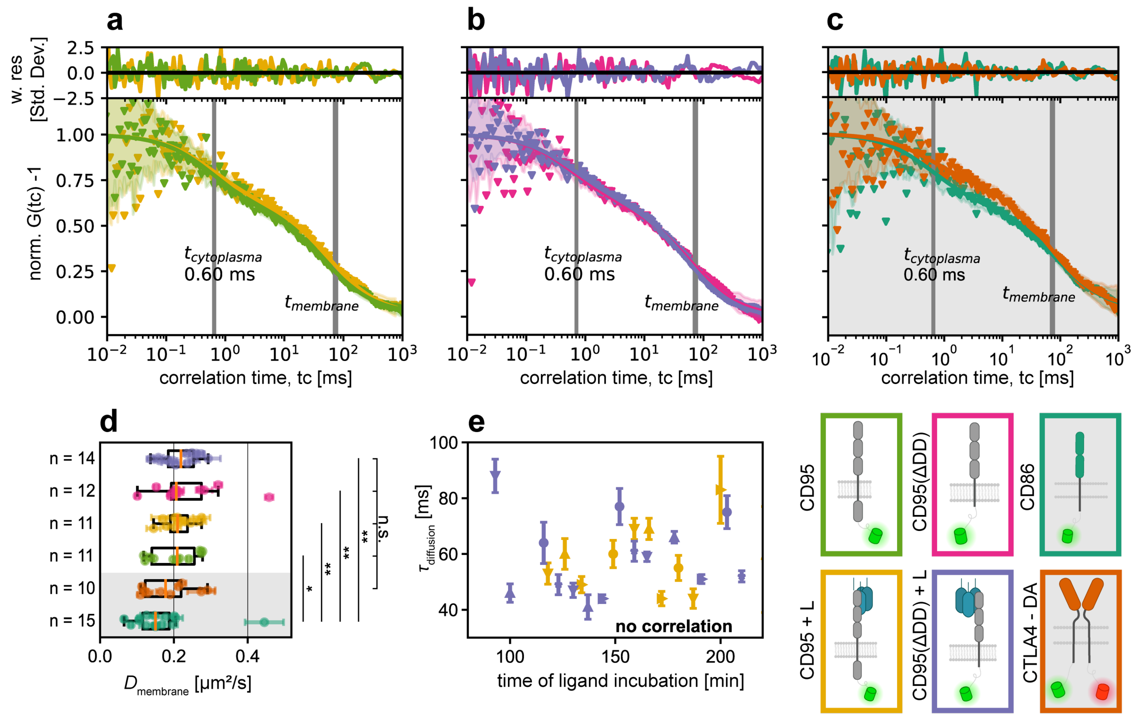

**Supplementary Figure 3: Live cell FCS to obtain diffusion times.**

Exemplary curves are shown for **a)** CD95 before (green) and 159 minutes after ligand addition (yellow) **b)** CD95(ΔDD) before (magenta) and 178 minutes after ligand addition (purple) **c)** CD86 monomeric control (teal) and CTLA4 dimer control (orange). All curves were fitted with two diffusion terms (compare methods and Equation (4)). The cytoplasmic diffusion term was fitted globally over 11 curves for the CD95 sample and fixed to this value for all other samples (see methods). **d)** Membrane diffusion constants were obtained from the membrane diffusion times. Mann-Whitney U-test was used to test for significance (*$p < 0.05$, **$p < 0.01$). **e)** Membrane diffusion time plotted against time since Ligand addition. No significant change was observed. At least ten different positions from at least 7 different cells were measured per sample and fitted subsequently. Legend: schematic representation of the samples used. Cytoplasmic and membrane fractions reported in Supplementary Table 1.

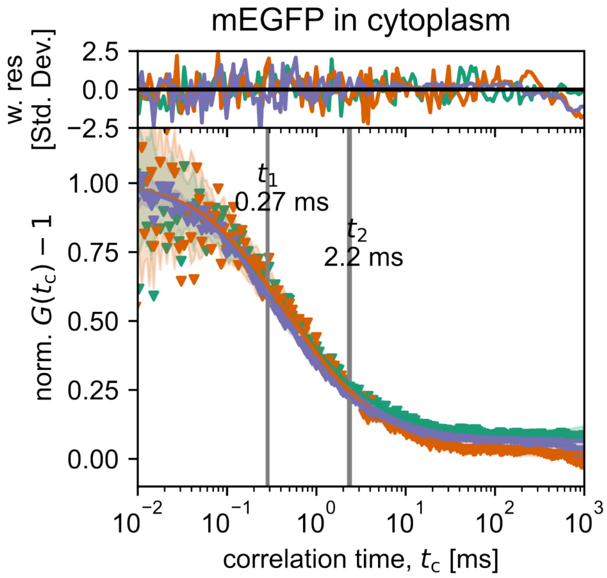

**Supplementary Figure 4: FCS curves of free mEGFP in cytoplasm.**
Free mEGFP in cytoplasm was fitted globally with two diffusion terms (Equation (4)), with a weighted average of 0.5 ms.

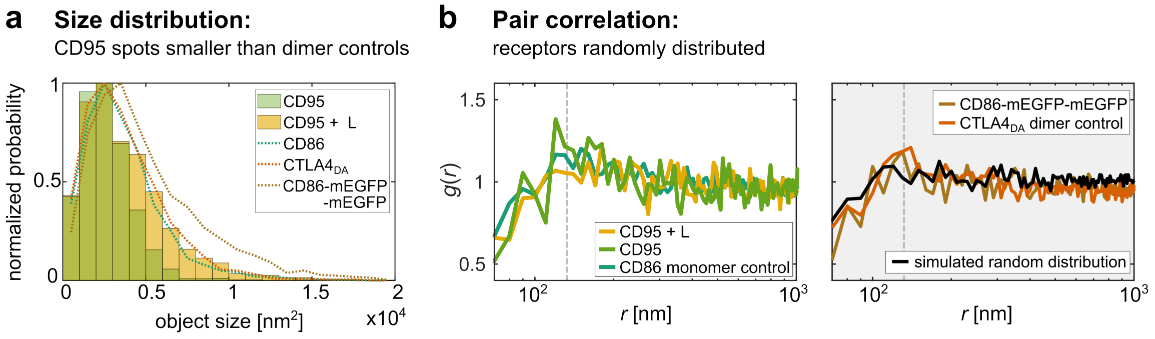

**Supplementary Figure 5: STED spot analysis of CD95  variant.**

**a)** Similar as for CD95(ΔDD) (Figure 3c), the distribution of full-length CD95 spot sizes (+/- ligand) is in the regime of monomer spot sizes. The slightly reduced size distribution of CD95 (no ligand) for larger spot sizes is correlating with a slightly lower expression level of the observed cells. The pseudo-dimer CD86-mEGFP-mEGFP show a slight shift to larger spot sizes. **b)** The pair correlation $g(r)$ (Equation (3)) of CD95 and dimer controls also shows a random distribution: Distances $r > 130$ nm (right of dashed line) with $g(r) \approx 1$ indicate a random distribution. A decrease in correlation for $r < 130$ nm arises from finite PSF size effects and not a particular distribution, as verified by simulations of randomly distributed spots with PSF (black curve, see *STED imaging and analysis* methods section).

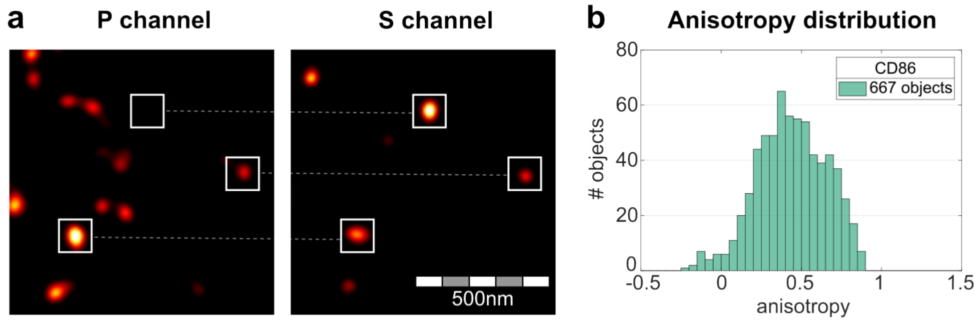

**Supplementary Figure 6: Polarization effect of STED samples.**
**a**) Deconvolved STED images of Hela CD95$^{KO}$ CD86-mEGFP stained with Atto647N α-GFP nanobody in parallel (P) and perpendicular (S) channel. The comparison of both images shows, that the emission of different spots is not equally distributed to both channels. **b)** The histogram shows the measured anisotropy of $n$ = 667 spots / objects (compare methods section and Equation (2) for details.). The spread in anisotropy confirms a strong polarization effect explaining the large spread in object intensities, even within one image.

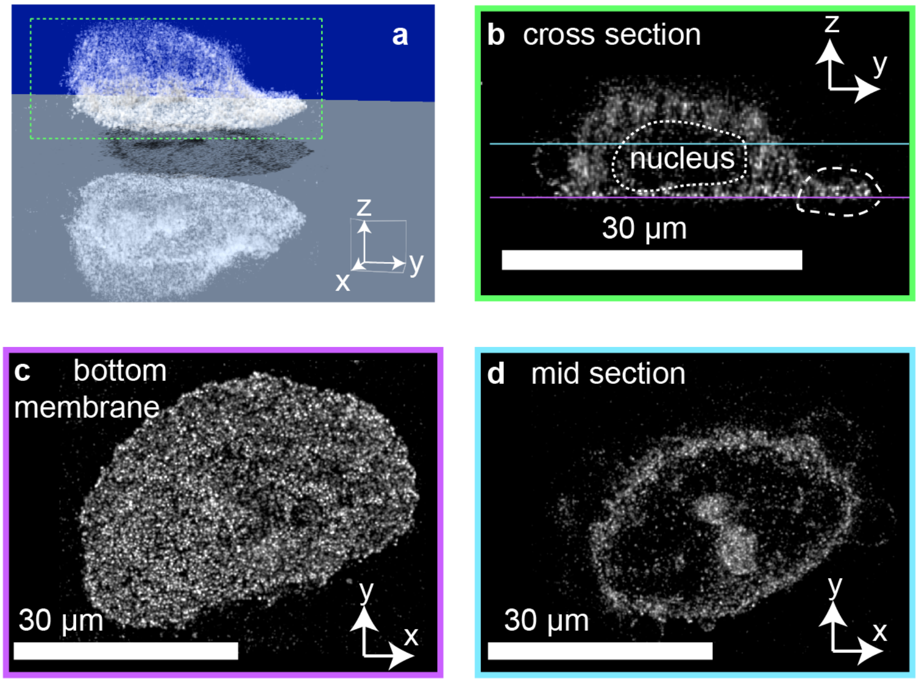

s

**Supplementary Figure 7: 3D confocal image of CD95 transfected fixed cell.**
**a)** Simulated fluorescence projection of 3D cell showing shadows on bottom plane. Cell was expressing mEGFP weakly and looks deflated due to the mounting process. **b)** xy cross section as indicated by dashed box in a. Dash-dotted line indicates the area where two membranes are in close proximity. Contour of nucleus is also shown. **c-d)** bottom and mid sections corresponding to colored planes in b. Figure highlights the need to measure cPBSA data below the nucleus to avoid having two membranes withing the confocal volume.

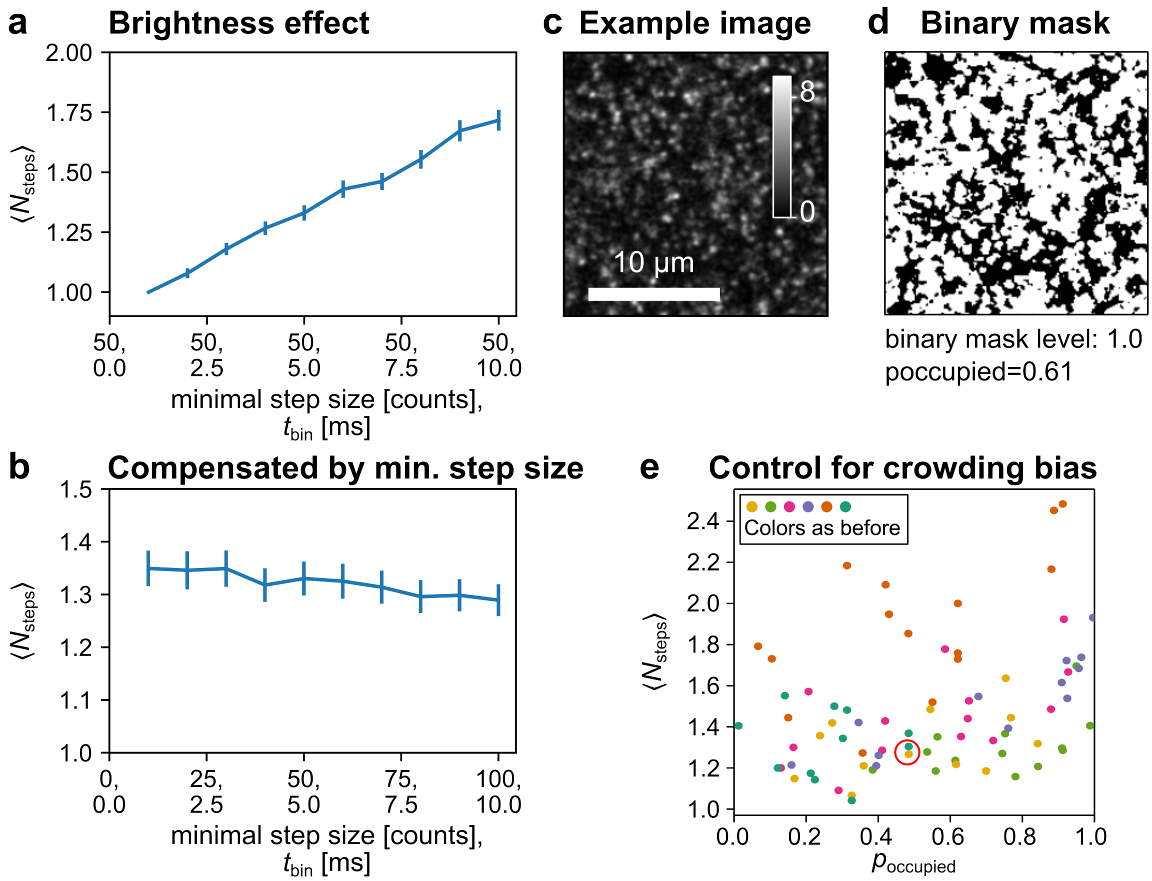

**Supplementary Figure 8: Controls for Confocal Photo-Bleaching Step Analysis.**
**a)** Effect of the brightness (effectively changed via $t_{\text{bin}}$) on cPBSA analysis on the average number of fluorophores for a CD86 dataset. **b)** Effect disappears when the threshold is increased in the same proportion as the $t_{\text{bin}}$. Data is obtained from the same dataset. **c)** Exemplary overview image smoothed with a 1 pixel sigma Gaussian filter from CD95 sample. **d)** Corresponding binary image illustrating $p_{\text{occupied}}$ as an indicator of multi-molecule events due to crowding. $p_{\text{occupied}}$ is determined as area fraction exceeding a signal intensity threshold of 1 pixel. **e)** To control for multi-step traces originating from multiple monomers or oligomers proximal in the confocal volume, the average numbers of bleaching steps $\langle N_{\text{steps}} \rangle$ according to cPBSA of one area is plotted against $p_{\text{occupied}}$, for that area. A weak positive correlation between $\langle N_{\text{steps}} \rangle$ and $p_{\text{occupied}}$ is visible as expected. As the spread of the occupancy probability was similar over all samples, this created no systematic shift in the data, wherefore no additional correction had to be introduced. Colors indicate samples matching the main text. Red circle corresponds to the area shown in c-d.

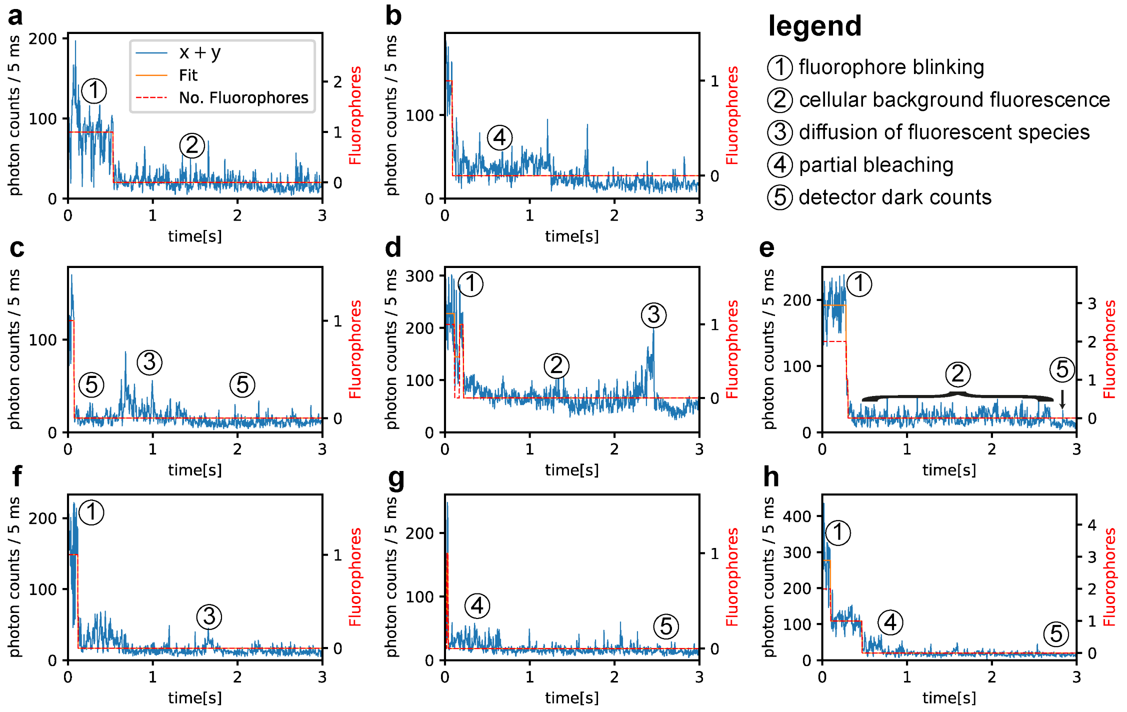

**Supplementary Figure 9: Exemplary traces for confocal photo bleaching step analysis.** **a-h)** Total fluorescent signal and fitted step trace, sources of noise are annotated. The most prominent sources of noise have been labelled in each graph for illustration purposes, although other sources are generally also present. **c-e)** Correlation curves of traces are plotted in Supplementary Figure 10, panels b-d, respectively. **h)** Two-step bleaching event show variation in step size. Traces were selected to illustrate noise sources and illustrate the overall data quality with little bias.

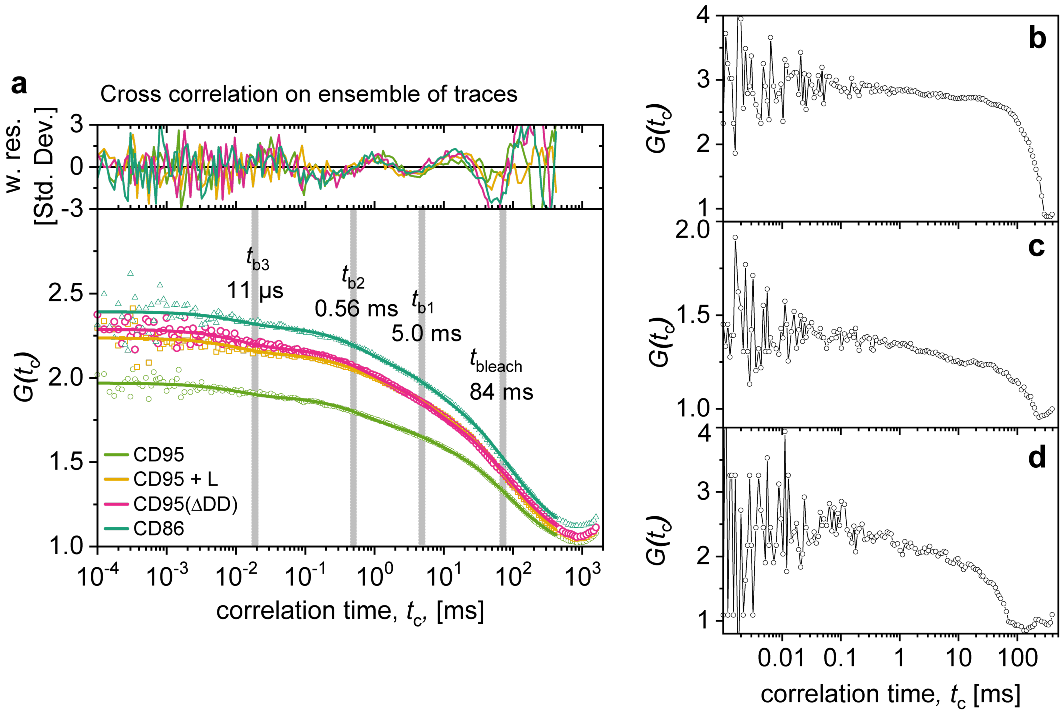

**Supplementary Figure 10: Correlation analysis on photo bleaching traces.**

**a)** Cross correlation curves of x- and y-polarization signals under circular polarization calculated for the ensemble of traces for each sample. Colors match the mainline figures. Curves were fitted with 3 bunching terms and 1 3D diffusion term as an approximation to model the bleaching behavior (see Equation (8)). As the bleaching statistics do not follow 3D diffusion statistics, we see a correlation in the residuals. All time parameters were fitted globally, whereas the fractions were left free (see Supplementary Table 4). We may obtain the parameters $\alpha$ and $k_{\text{on}}$ from the bunching fraction and bunching times, respectively, and predict the variance of trace sections (see Supplementary Note 3). Note that the bunching fractions were similar over different samples, indicating that the photophysical properties of mEGFP over different samples were similar. **b-d)** CD86 correlation curves correspond to traces c-e respectively of Supplementary Figure 9. As all photons correlate, a high signal correlation curve can be generated based on only a few photons, allowing single-molecule based correlation fits. Whereas the correlation due to blinking looks similar to the ensemble fits, the bleaching correlation varies per molecular assembly, as also the bleaching time and amplitude varies.

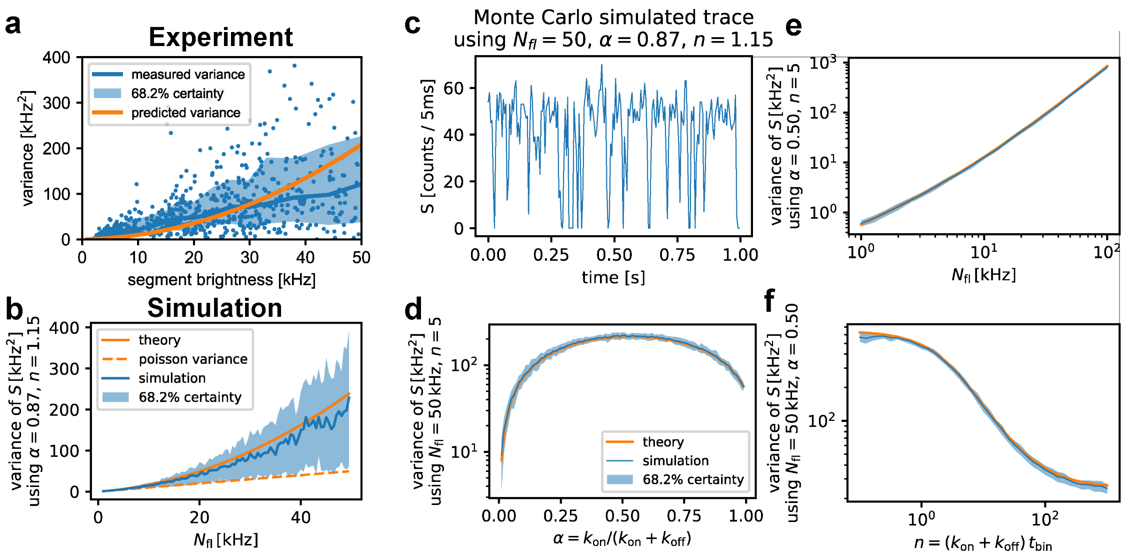

**Supplementary Figure 11: A single dark state predicts trace variance.**
A theoretical formula for the variance of an emitter under a single dark state (see Supplementary Note 3) is compared to measured and simulated trace variances. **a)** For each trace segment from CD86 traces measured on 22 July 2021 the segment intensity and variance was obtained from PBSA analysis (blue points). For each 50 consecutive intensities the median (blue line) and 15.9% - 84.1% quantiles (blue area) are determined to aid visualization. Supplementary Equation (S7) is used to predict the variance for each segment using $\alpha$ and $n$ obtained from FCS[12] and $N_{\mathrm{fl}}$ for each segment. **b)** Monte Carlo traces using identical $\alpha, n$ , trace duration as in a) match the theoretical prediction. The short trace duration (84 ms, see Supplementary Figure 10) causes a large spread in the variance due to the varying number of blinks in this period, matching the spread observed in a). For reference the variance based on pure Poisson noise is shown. **c)** Exemplary Monte Carlo trace over 1 second under conditions typical to mEGFP measurements. **d-f)** Similar to b, but using a simulation time of 1 second and scanning the whole parameter space. Legend for all as given in d).

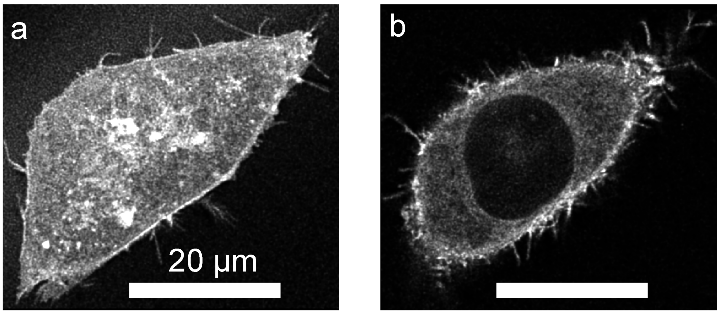

**Supplementary Figure 12: CD95 transfected cells imaged by confocal microscopy.**
**a)** Live cell transfected with CD95 bottom membrane of 3D stack. Image was recorded according to Nyquist sampling and deconvolved using Express Deconvolution Huygens Professional 21.10.0, SNR 6. **b)** mid-segment of same 3D, color scale same as in a. Signal is brightest in the membrane, but also cytoplasmic intensity is visible. No Fluorescent signal is present in the nucleus.

### References

1 B.V., S. V. I. *Backprojected Pinhole Calculator*, <https://svi.nl/Olympus_FV1000> (2021).

2 Wohland, T., Rigler, R. & Vogel, H. The standard deviation in fluorescence correlation spectroscopy. *Biophysical Journal* **80**, 2987-2999 (2001). https://doi.org:10.1016/s0006-3495(01)76264-9

3 van der Voort, N. T. M., Bartels, N., Monzel, C. & Seidel, C. A. M. Quantifying the Spatio-temporal Evolution of Protein Interactions using Cell Lifetime FRET Image Spectroscopy (CELFIS). *bioRxiv* (2022).

4 Koppel, D. E. Statistical Accuracy in Fluorescence Correlation Spectroscopy. *Physical Review A* **10**, 1938-1945 (1974). https://doi.org:10.1103/PhysRevA.10.1938

5 Eggeling, C. *et al.* Data registration and selective single-molecule analysis using multi-parameter fluorescence detection. *Journal of Biotechnology* **86**, 163-180 (2001). https://doi.org:10.1016/s0168-1656(00)00412-0

6 Oracz, J., Westphal, V., Radzewicz, C., Sahl, S. J. & Hell, S. W. Photobleaching in STED nanoscopy and its dependence on the photon flux applied for reversible silencing of the fluorophore. *Scientific Reports* **7**, 11354 (2017). https://doi.org:10.1038/s41598-017-09902-x

7 Cranfill, P. J. *et al.* Quantitative assessment of fluorescent proteins. *Nature Methods* **13**, 557-562 (2016). https://doi.org:10.1038/nmeth.3891

8 Duan, C. X. *et al.* Structural Evidence for a Two-Regime Photobleaching Mechanism in a Reversibly Switchable Fluorescent Protein. *Journal of the American Chemical Society* **135**, 15841-15850 (2013). https://doi.org:10.1021/ja406860e

9 Potma, E. O. *et al.* Reduced protein diffusion rate by cytoskeleton in vegetative and polarized Dictyostelium cells. *Biophysical Journal* **81**, 2010-2019 (2001). https://doi.org:10.1016/s0006-3495(01)75851-1

10 Hennen, J., Hur, K. H., Saunders, C. A., Luxton, G. W. G. & Mueller, J. D. Quantitative Brightness Analysis of Protein Oligomerization in the Nuclear Envelope. *Biophysical Journal* **113**, 138-147 (2017). https://doi.org:10.1016/j.bpj.2017.05.044

11 Wenger, J. *et al.* Diffusion analysis within single nanometric apertures reveals the ultrafine cell membrane organization. *Biophysical Journal* **92**, 913-919 (2007). https://doi.org:10.1529/biophysj.106.096586

12 Barth, A. *et al.* Unraveling multi-state molecular dynamics in single-molecule FRET experiments. I. Theory of FRET-lines. *Journal of Chemical Physics* **156**, 141501 (2022). https://doi.org:10.1063/5.0089134
